# Variable expression of linguistic laws in ape gesture: a case study from chimpanzee sexual solicitation

**DOI:** 10.1101/2021.05.19.444810

**Authors:** Alexandra Safryghin, Catharine Cross, Brittany Fallon, Raphaela Heesen, Ramon Ferrer-i-Cancho, Catherine Hobaiter

## Abstract

Two language laws have been identified as consistent patterns shaping animal behaviour, both acting on the organisational level of communicative systems. Zipf’s law of brevity describes a negative relationship between behavioural length and frequency. Menzerath’s law defines a negative correlation between the number of behaviours in a sequence and average length of the behaviour composing it. Both laws have been linked with the information-theoretic principle of compression, which tends to minimise code length. We investigated their presence in a case study of male chimpanzee sexual solicitation gesture. We failed to find evidence supporting Zipf’s law of brevity, but solicitation gestures followed Menzerath’s law: longer sequences had shorter average gesture duration. Our results extend previous findings suggesting gesturing may be limited by individual energetic constraints. However, such patterns may only emerge in sufficiently-large datasets. Chimpanzee gestural repertoires do not appear to manifest a consistent principle of compression previously described in many other close-range systems of communication. Importantly, the same signallers and signals were previously shown to adhere to these laws in subsets of the repertoire when used in play; highlighting that, in addition to selection on the signal repertoire, ape gestural expression appears shaped by factors in the immediate socio-ecological context.

## Introduction

Over the past 100 years, important statistical regularities have been described across human languages and in other communicative systems such as genomes, proteins, and animal vocal and gestural communication (Altmann & Gerlach, 2016; Bentz & Ferrer-I-Cancho, 2016; Börstell et al., 2016; Hernández-Fernández et al., 2019; Köhler et al., 2005; Menzerath, 1954; Naranan & Balasubrahmanyan, 2000; Sanada, 2008; Semple et al., 2022; Wang & Chen, 2015; Zipf, 1936). These regularities are hypothesized to be manifestations of the information theoretic principle of compression (Ferrer-i-Cancho, Bentz, et al., 2022; Semple et al., 2022). Compression is a particular case of the principle of least effort (Zipf, 1949) – a principle that promotes the outcome that requires the least amount of energy to produce or achieve – and thereby promotes coding efficiency (Ferrer-i-Cancho et al., 2013). In communication, compression is expressed as a pressure towards reducing the energy needed to compose a code but limited by the need to retain the critical information in the transmission (Cover & Thomas, 2006; Ferrer-i-Cancho et al., 2022).

Among the statistical patterns predicted by compression at different levels of organization, Zipf’s law of brevity and Menzerath’s law have been at the centre of recent attention in studies of human and non-human communication. Zipf’s law of brevity is the tendency for more frequent words to be shorter in length (Strauss et al., 2007; Zipf, 1949), and is generalised as the tendency for more frequent elements of many kinds (*e*.*g*., syllables, words, calls) to be shorter or smaller (Ferrer-i-Cancho et al., 2013) – with similar patterns found at different levels of analysis, for example in speech at the level of words (Strauss et al., 2007) syllables (Rujević et al., 2021), and phonemes (Hernández-Fernández et al., 2019). As well as being found in human spoken, signed, and written languages (Bentz & Ferrer-I-Cancho, 2016; Börstell et al., 2016; Hernández-Fernández et al., 2019; Sanada, 2008; Wang & Chen, 2015), Zipf’s law of brevity has been identified in the short-range communication of diverse taxa: dolphins (Ferrer-i-Cancho et al., 2022), bats (Luo et al., 2013), penguins (Favaro et al., 2020), hyraxes (Demartsev et al., 2019), and various primates (macaques: Semple et al., 2013; marmosets: Ferrer-i-Cancho & Hernández-Fernández, 2013; gibbons: Huang et al., 2020; Indri indri: Valente et al., 2021), as well as in genomes (Naranan & Balasubrahmanyan, 2000).

At the level of constructs, Menzerath’s law states that “*the greater the whole, the smaller its constituents*” (Altmann, 1980; Köhler, 2012; Menzerath, 1954); for example: longer sentences have words of shorter average length, and words with more syllables contain syllables of shorter length. Menzerath’s law (and its mathematical expression known as the Menzerath-Altman’s law) has been identified in human spoken and signed languages (Altmann, 1980; Andres et al., 2021), genomes (Ferrer-i-Cancho & Forns, 2009; Li, 2012), music (Boroda & Altmann, 1991), and in the communication of dolphins (Ferrer-i-Cancho et al., 2022), penguins (Favaro et al., 2020), and primates (geladas: Gustison et al., 2016; chimpanzees: Fedurek et al., 2017; Heesen et al., 2019; gibbons: Clink et al., 2020; Huang et al., 2020; gorillas: Watson et al., 2020; Indri indri: Valente et al., 2021;). While many studies focused on vocal communication, several have now explored these statistical regularities in gestural and signed domains. For example, the use of Swedish Sign Language in (semi-)spontaneous conversation was found to follow a pattern of more frequently used signs being shorter in duration (Börstell et al., 2016). Zipf’s law of brevity was also found in fingerspelling, with a negative relationship between mean fingerspelled sign duration and frequency (Börstell et al., 2016). Similarly, Czech sign language was found to follow Menzerath’s law (Andres et al., 2021). Work in non-human gesture has, to date, been more focused on context-specific signal usage, for example: Zipf’s law of brevity was found in the surface behaviour of dolphins (such as tail-slapping; Ferrer-i-Cancho & Lusseau, 2009) but not in the overall repertoire of play gestures of chimpanzees, where it was only present in subsets, although these gestures did follow Menzerath’s law (Heesen et al., 2019).

Chimpanzee gestural communication represents a powerful non-human model in which to explore compression and language laws. Apes have large repertoires of over 70 distinct gesture types (Byrne et al., 2017); as compared to vocal communication, gestural repertoires are larger and are more flexibly deployed, with individual gesture types used to achieve multiple goals (Bard et al., 2019; Call & Tomasello, 2007; Hobaiter & Byrne, 2011a; Liebal et al., 2004). Gestures are also used intentionally, *i*.*e*., to reach social goals by influencing the receivers’ behaviour or understanding (Graham et al., 2018; Hobaiter & Byrne, 2011a, 2014; Schel et al., 2013), and flexibly across contexts (Call & Tomasello, 2007; Hobaiter & Byrne, 2011a; Liebal et al., 2004). Nevertheless, Heesen et al.’s (2019) results support an increasingly diverse range of findings that show variation in the extent and expression of language laws, suggesting that while they appear statistically universal there is room for exceptions and/or variation in patterning at different levels of the communicative construct (Semple et al., 2022).

Although a lack of evidence supporting Zipf’s law of brevity has been previously reported (e.g., European heraldry: Miton & Morin, 2019; computer-based neural-networks: Chaabouni et al., 2019), these remain rare exceptions, and in non-human animal communication have typically only been reported in long-distance vocal communication (*e*.*g*. gibbon song: Clink et al., 2020; bats: Luo et al., 2013; although *cf*. female hyrax calls: Demartsev et al., 2019) where the impact of distance on signal transmission fidelity may have a particularly strong effect on the costs of compression (Ferrer-i-Cancho et al., 2013; Gustison et al., 2016; Semple et al., 2022). Thus, at present, the repertoire-level absence of Zipf’s law of brevity in chimpanzee gesture remains a conundrum.

One explanation for a repertoire-level absence of Zipf’s law of brevity – as seen in some long-distance signals – is that the context in which signals are produced may impact the emergence and expression of these patterns. Specifically, in the case of chimpanzee gestures, the absence of a pattern resembling Zipf’s law of brevity may result from the use of gestures produced during play. Expressions of linguistic laws in biological systems reflect pressures that shape efficient energy expenditure (Semple et al., 2022). Play is produced when there is an excess of time and energy (Held & Špinka, 2011; Pellis & Pellis, 1996; Smith, 2014), thus, the energetic need to reduce signal effort through increased compression may be limited. As a result, it remains unclear whether the failure of Zipf’s law of brevity in chimpanzee gesture was due to the use of gestures from within play, or whether it reflects a system-wide characteristic.

In both signed languages and human gesturing, distinctions are made between different components of their production. First there is the *preparation* of the signal, then the *action stroke* which represents the movement that defines the gesture as of a particular type; an individual can then choose to further *hold* the stroke or repeat it, until they decide to stop gesturing and return the limb to rest during *recovery* from the gestural action (Kendon, 2004). For example: in a reach gesture this would correspond to the movement of the hand into position (*preparation*), the extension of the arm and hand towards the recipient (*action stroke*), the (optional) maintenance of the extension (*hold*), and finally the return of the hand and arm to a resting state (*recovery*). All four of these phases require some energetic investment to produce, but there may be variation across them, and aspects such as preparation and recovery may be nearly, or entirely, absent where several gestures are strung together. In some gesture types, their production does not include a *hold* phase (e.g., hit, jump, throw object); we term these *fixed* duration gestures, as the duration of their expression is relatively constrained across instances of production. Other gesture types can include a *hold* phase (for example: reach, object shake, swing) which may or may not be present, and, where present, may vary substantially in length; we term these *loose* duration gestures. There may be differences in the emergence of Zipf’s and Menzerath’s laws regarding the different components of gesture production. Menzerath’s law acts from a proximate perspective on the building of communicative sequences in a specific communicative instance: for example, gestures produced in longer sequences may be shortened by variation the duration of components such as the shortening of the *hold* phase in loose gesture types. In contrast, Zipf’s law acts on gesture types across instances of use – and as such may be less sensitive to the immediate context of production.

Another possible explanation for the variation in the emergency of compression in ape gesture is that the ability to detect linguistics laws, particularly where they are only subtly expressed, appears to require powerful datasets. The exploration of statistical patterns in human languages often employs corpora containing millions of data points (e.g., Hatzigeorgiu et al., 2001). In contrast, in ape gesture, as in many studies of non-human communication, datasets are substantially smaller (in the thousands). In chimpanzee play, the large repertoire expressed limits the frequency with which particular gesture types are represented.

We address this open question in a case study of chimpanzee gestural communication in sexual solicitation. While gesture is relatively under-studied in this area, sexual solicitations have been contrasted with early descriptions of gesture from studies of captive ape play, as an example of gesture in a relatively more evolutionarily or biologically ‘relevant’ context for communication (in terms of associated risks and/or impact on reproduction) (Hobaiter & Byrne, 2012; *c*.*f*. Call & Tomasello, 2007). Chimpanzees, particularly male chimpanzees, employ prolific use of individual gestures and gesture sequences in sexual solicitations. As solicitations are often vigorous, chimpanzees incorporate regular use of gesture types that include both visual and audible information (Hobaiter & Byrne, 2012; Nishida, 1980). While a range of gesture types are employed, these are typically a smaller sub-set of the available repertoire – *c*.*f*. play where the majority of gesture types are deployed. Successful gestures can lead directly to sexual behaviour, such as inspection or copulation, as well as to a consortship, in which the female follows the male away from other individuals in the group so that he maintains exclusive sexual access (Tutin, 1979). Both direct solicitation and consortship and are key strategies for individual fitness (Tutin, 1979; Watts, 2015), and as such behaviour associated with them is likely subject to strong selective pressures. The energetic costs of lactation mean that adult female chimpanzees typically concieve only once every 4-5 years (Clark, 1977; Thompson, 2013). So while there are typically 60-80 individuals in a group, the operational sex ratio of available females in estrus may be very small, and males show substantial variation in reproductive success (Newton-Fisher et al., 2009; Tutin, 1979). Although highly important, the performance of sexual solicitations may come with significant costs: besides the energetic expenditure in producing these signals, there is a risk of potentially aggressive competition both from other males in their own community (Fawcett & Muhumuza, 2000; Tutin, 1979) as well as potentially lethal attacks from males in neighbouring groups (Wilson et al., 2014). For example, during consortships individuals may travel to the boundaries of their home area, increasing the risk of encounters with neighbouring individuals. Thus, there are substantial advantages to avoiding potential eavesdroppers within, and particularly outside of, one’s community (Hobaiter et al., 2017). Therefore, on one hand individuals benefit from producing conspicuous energetic signals to attract females, often having to insist to secure mating; on the other, the production of highly conspicuous signals should be compressed to reduce the risks associated with competition from both within and outside the group.

To assess compression in the sexual solicitation gestures of wild male chimpanzees, we tested for patterns predicted by Zipf’s law of brevity and Menzerath’s law, both at the level of single gesture types and gesture sequences, respectively. To investigate Zipf’s law of brevity and Menzerath’s law we fitted two generalised linear mixed models. The first model explored the presence of Zipf’s law assigning gesture duration as the response variable, proportion of gestures within the dataset and category of gesture (manual vs whole body) as fixed factors, and signaller’s ID, sequence ID, and gesture type as random factors. The second model tested for Menzerath’s law and had gesture duration as response variable, sequence size as a fixed factor, and proportion of whole-body gestures in the sequence (PWB), signaller ID, sequence ID, and gesture type as random factors. We included information on the category of the gesture to allow for comparisons with human studies, in which gestures are mostly manual. We provide matched models that describe the patterns of expression both across (i) all males in our data, (ii) for a single prolific individual and (iii) for the remaining individuals. In doing so, we provide an initial assessment of the distribution of our findings across male chimpanzee gesturing in this context and provide an expanded assessment of compression in ape gestural communication.

## Results

We measured *N*=560 sexual solicitation gestures from 173 videos of 16 wild, habituated male East African chimpanzees (*Pan troglodytes schweinfurthii*) gesturing to 26 females. Within the 560 gestural instances (from now *tokens*), we identified 26 gesture types: 21 manual gestures and 5 whole-body gestures (Figure 1; for definitions for full repertoire definitions see Table S1 in supporting Information 1) performed by 16 male chimpanzees aged 10-42 years old. On average, each individual produced a median of 11.5 ± 70.7 gesture tokens (range 2-290). One male, Duane, was particularly prolific (*n*=290 gesture tokens; other males 2-76). To provide context as to what extent our findings are generalizable, we provide matched analyses using both the full dataset and the dataset limited to Duane only. An analysis of the data excluding Duane is available in the supplementary information.

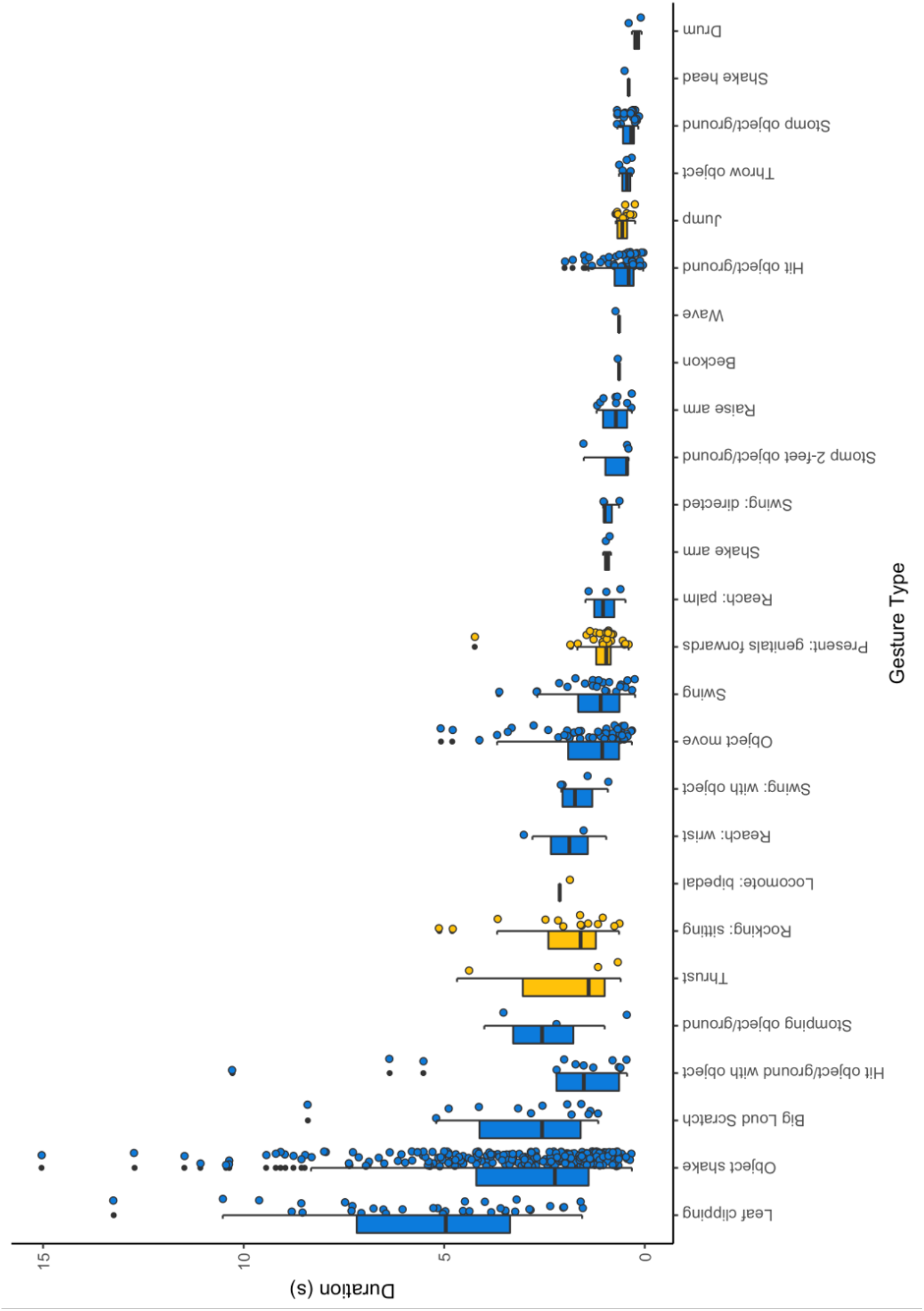

Gesture token duration was measured via analysis of video data with a minimum unit of 0.04s (one frame). Duration ranged from 0.04-15.04 seconds (median: 1.56 ±2.35s). If consecutive gesture tokens were performed with less than 1s in between them, they were considered to form a sequence (Heesen et al., 2019; Hobaiter & Byrne, 2011b). We detected a total of 377 sequences, with each male performing a median of 8 ±44.54 sequences (range 1-181 sequences). Sequence length ranged from 1 to 6 tokens (Table 1). For analyses of Menzerath’s law we excluded 18 sequences for which we were unable to identify the duration of all the consecutive gesture tokens performed, resulting in the analysis of 359 sequences, containing a total of 530 gesture tokens. 244 sequences were composed of a single token, the remaining 115 sequences had length *n*>1. Of the 115 sequences analysed that were composed of 2 or more gesture tokens; 26 (23%) were formed by the repetition of the same gesture type, whereas the remaining 89 (77%) included more than one gesture type (Table 1).

**Table 1.**
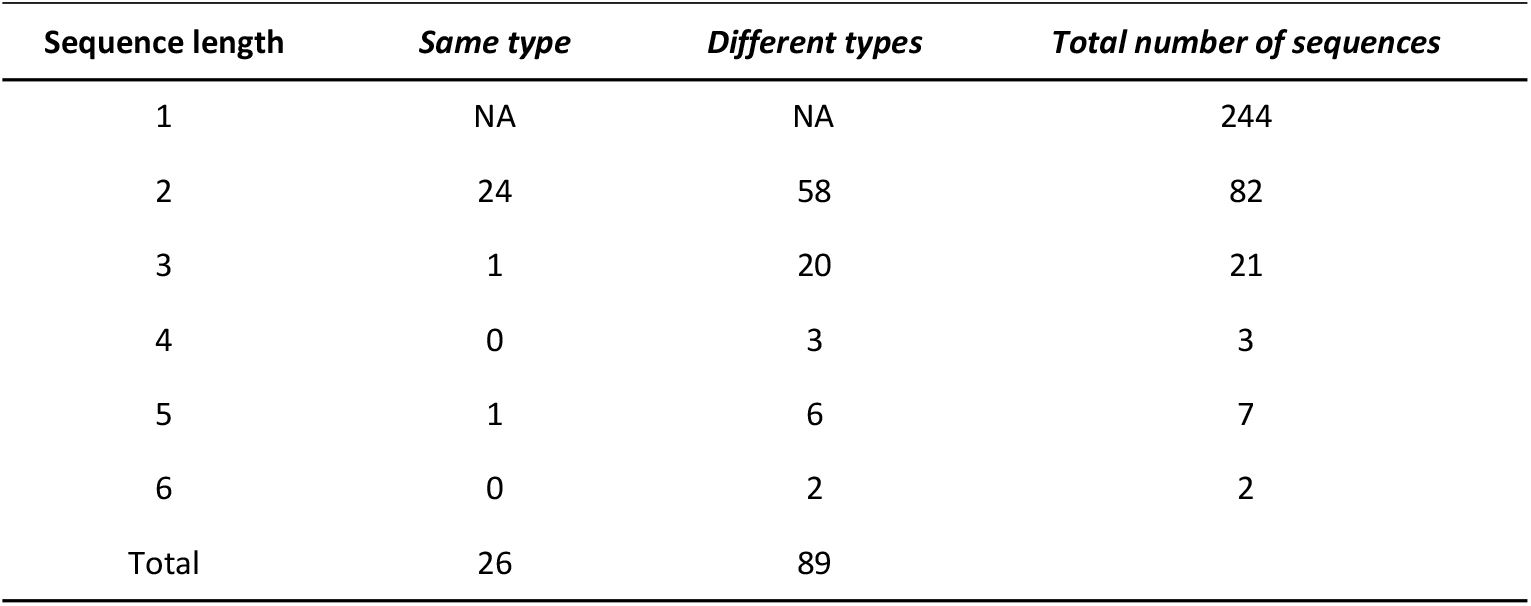

| **Sequence length** | ***Same type*** | ***Different types*** | ***Total number of sequences*** |
| --- | --- | --- | --- |
| 1 | NA | NA | 244 |
| 2 | 24 | 58 | 82 |
| 3 | 1 | 20 | 21 |
| 4 | 0 | 3 | 3 |
| 5 | 1 | 6 | 7 |
| 6 | 0 | 2 | 2 |
| Total | 26 | 89 |  |

### Do chimpanzee sexual solicitation gestures follow Zipf’s law of brevity?

To test for Zipf’s law we ran a Bayesian generalised linear model (Zipf-model), with the log of gesture duration as the response variable and the proportion of gesture type within the dataset as a fixed factor (see Supporting information 2 for further detail). The gesture duration data was log-transformed following an analysis of data distribution. We included category of gesture as a control, and signaller ID, sequence ID, and gesture type as random factors. The Zipf-model fitted the data better than a null model that did not include the proportion of gesture type as a fixed effect (Leave-one-out [LOO] difference and s.d.= -0.7 ± 0.3). For Zipf-model effects Bulk ESS and Tail ESS were >100 and 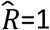. However, the proportion of gesture type did not have a substantial effect on the duration of gestures (Supporting information 3, Table S5; b = 0.90, s.d. = 1.26, 95% Credible Intervals (CrI) [-1.25, 3.81], Figure 2A). When testing the subset of data containing only the gestures produced by Duane, the full model and null model testing for Zipf’s law showed similar fit (LOO difference: -0.1 ± 0.7; Supporting information 3, Table S6, Figure 2B). Similarly, in the same analysis on data from all individuals except Duane, the full model was no different from the null model (LOO difference: -0.5±0.5; Supporting information 3, Table S7).

**Figure 2.**
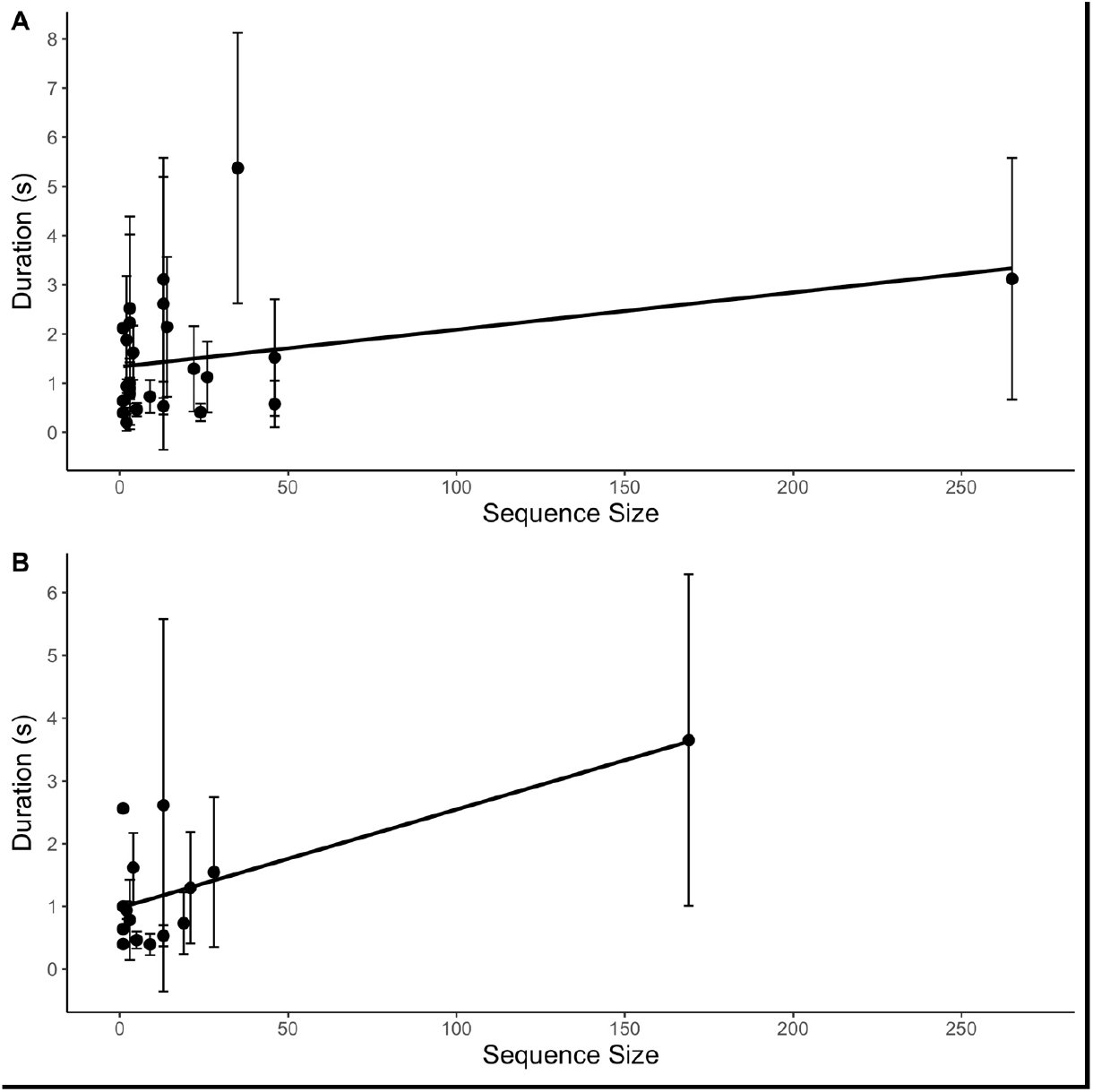
Relationship between frequency of occurrence and gesture duration for the full dataset (A) and Duane only data (B). Points represent the mean duration of each gesture type, with error bars showing the standard deviation from the mean. Black line indicates regression slope. 159×159mm (220 × 220 DPI)

### Do chimpanzee sexual solicitation gesture sequences follow Menzerath’s law?

To test for Menzerath’s law we ran a second Bayesian model (Menzerath-model) with the log of the gesture duration as response variable, the sequence size as fixed factor, the proportion of whole-body gestures within the sequence (PWB) as a control, and the signaller ID and sequence ID as random factors. The Menzerath-model fitted the data better than the null model (LOO difference: -7.7 ± 4.1). All predictors had Bulk ESS and Tail ESS>100 as well as 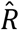 values =1. Sequence size had a substantial negative effect on gesture duration within sequence (Supporting information 3, Table S8; b = -0.18, s.d. = 0.04, 95% CrI [-0.26, -0.11]; Figure 3A). Similar results were found when running the same Menzerath-model but limited to gestures produced by Duane: the full model fitted the data better than the null (LOO difference: -13.5 ± 4.5), all predictors had Bulk ESS and Tail ESS>100, 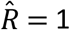 and sequence size had a substantial negative effect on gesture duration (Supporting information 3, Table S9, Figure 3B; b = -0.23, s.d. = 0.04, 95% CrI [-0.31, -0.15]). In contrast, where Duane’s data were excluded, the full model was similar to the null model, suggesting no clear pattern consistent with Menzerath’s law (LOO difference: -0.3±0.8; Supporting information 3; Table S10). Visual inspection of the data plotted per individual suggests that detection of a pattern consistent with Menzerath’s law may be impacted by sample size (Supporting Information 4).

**Figure 3.**
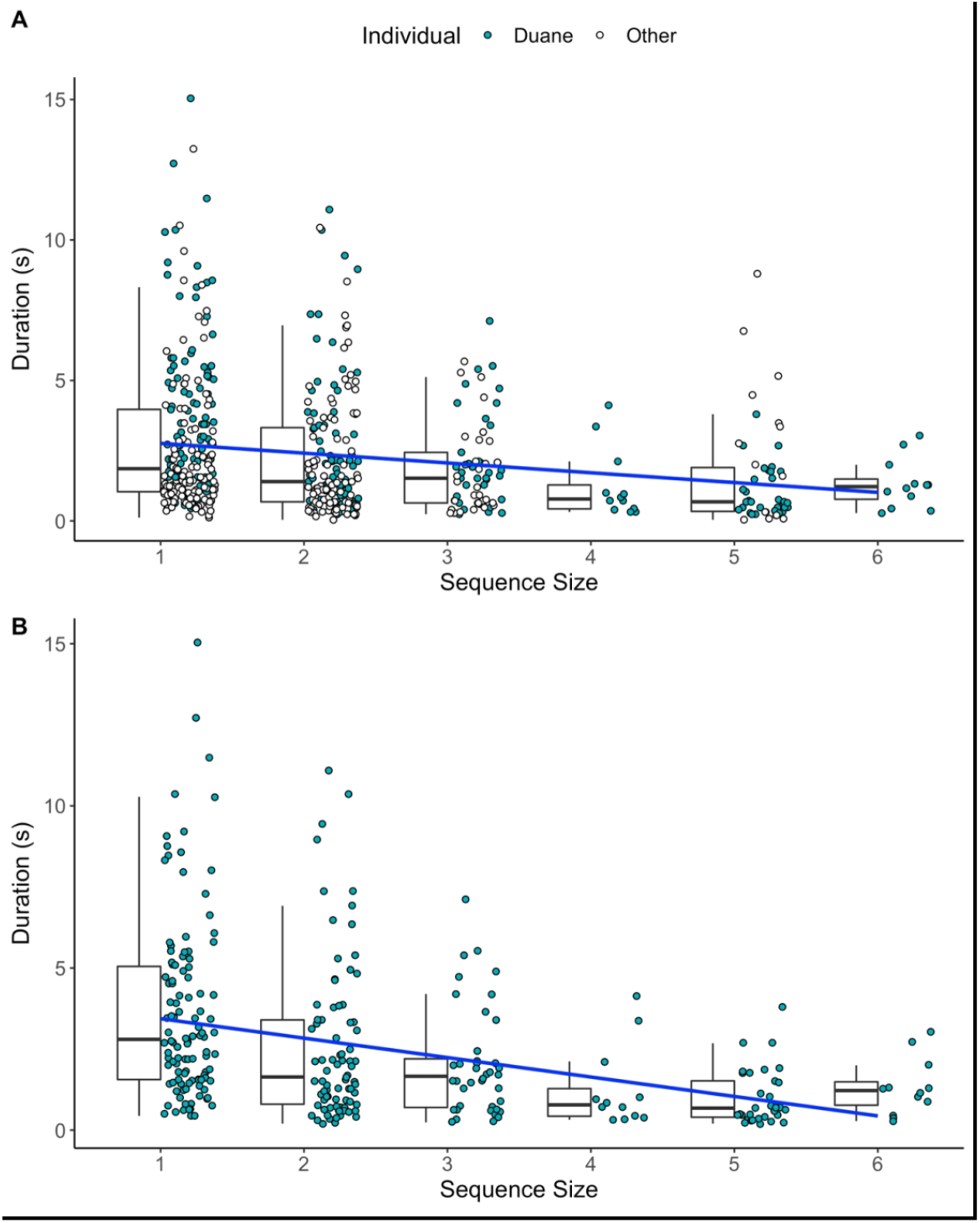
Relationship between sequence size and gesture duration for the full dataset (A) and Duane only data (B). Boxplots represent the median (black bar), the interquartile range – IQR (boxes), and maximum and minimum values excluding outliers (whiskers). Points represent individual gesture tokens, ordered by the length of the sequence they were performed in. Gestural tokens belonging to the individual Duane are indicated in light blue. White circles indicate gesture tokens belonging to all other individuals. Blue line indicates regression slope. 159×199mm (220 × 220 DPI)

We note that the sample size of sequences of four tokens or longer is smaller than those of one to three tokens (Table 1), which may have contributed to the apparent tailing off of a clear relationship in Figure 3A and Figure 3B. In addition, longer sequences were formed of a) a mix of *loose* and *fixed* duration gestures or b) only *loose* duration gestures (see supporting information 5, Figures S5 and S6). Thus, the emergence of Menzerath’s law could not be explained by a shift in preference from *fixed* to *loose* gestures with increasing sequence length.

## Discussion

Chimpanzee sexual solicitation gestures did not follow Zipf’s law of brevity: the frequency of gesture type within the dataset did not predict gesture duration in any of our samples.

However, sequences of chimpanzee solicitation gestures did follow Menzerath’s law: longer sequences of gestures were made up of gestures of shorter average length. Our dataset was limited both by its relatively small size (*c*.*f*. Heesen et al., 2019 on chimpanzee play gestures) and in its bias towards a single highly prolific individual (Duane). As a result, we consider it a case-study; however, the pattern was present in both the Duane’s data and in the full dataset, as well as in a range of alternative analyses (Supporting Information 6). In the reduced dataset excluding Duane we did not find a pattern consistent with Menzerath’s law; however, detection of the pattern may have been limited by the small sample size available in the remaining data set.

These results represent a further absence of evidence in support of Zipf’s law of brevity in great ape gestural communication (Heesen et al., 2019) and support the wider finding that – unlike most other close-range systems of communication described to date – the expression of pressure for compression and efficiency may be variably expressed in ape gesture (Börstell et al., 2016; Ferrer-i-Cancho et al., 2013; Semple et al., 2022). It particularly highlights that compression does not act on communicative systems uniformly: 20 of the 26 gesture types described here as used in sexual solicitations overlapped with those used in play (Heesen et al., 2019). Data were collected from the same community over the same period, and although both studies provided a null result when analysing the full gestural repertoire, Zipf’s law was found in subsets of the play gestures but not in the gestures when used in sexual solicitations. Moreover, when running traditional correlation analyses in which features such as signaller identity, or gesture type could not be controlled for, we found a tendency for an opposite Zipf’s law pattern – particularly in manual gestures (Supporting Information 6). Visual inspection of the Figure 3 shows the substantial variation in the duration of gestures across instances of communication, as well as an apparent decrease in a clear relationship between gesture duration and sequence size where sample size was small (such as for longer sequences). Together these findings suggest that the expression of these laws is nuanced by aspects of the communicative landscape in which they are deployed, and that large samples may be needed to detect sometimes subtle relationships. Future work could specifically explore variation in the detection of these patterns at different sample sizes, for example by randomised subsetting of sufficiently large datasets. As Semple et al. (2022) suggest, apparent ‘failures’ may be of substantial assistance in exploring the boundaries of the theoretical framework of these laws, helping to define the characteristics that shape both their emergence and variation in their expression.

In contrast to vocal communication across primate species, in chimpanzee sexual solicitations ‘inefficiency’ in signalling effort by the signaller appears to be at times slightly favoured. However, these gestures appear to remain effective in terms of achieving the signaller’s goal of successful communication in a context vital for reproductive success. Given the long inter-birth intervals and active mate guarding (Muller & Wrangham, 2009), chimpanzee paternity is often heavily biased towards higher-ranking individuals (Newton-Fisher et al., 2009). With so few opportunities to mate, sexual solicitations may represent one of the most evolutionarily important contexts in which chimpanzee gestures are produced. Where the costs of signal failure are high, there is a pressure against compression and towards redundancy, as in chimpanzees’ use of gesture-vocal signal combinations in agonistic social interactions (Hobaiter et al., 2017). While there are examples of vocal communication systems used in biologically ‘relevant’ contexts that adhere to Zipf’s brevity law (Favaro et al., 2020), the benefits of successful communication to individual fitness in chimpanzee solicitation appear to outweigh the energetic costs associated with the production of a vigorous and conspicuous signal. Nevertheless, given that we see a relatively consistent expression of Menzerath’s law across gesture use in sexual solicitation as in play, even the production these prolonged and conspicuous signals appear to remain constrained by physiological mechanisms of gestural production. As for primate vocal communication (Fedurek et al., 2017; Gustison et al., 2016), where breathing constraints and energetic demands of vocal production were considered drivers for the emergence of Menzerath’s law patterns, increased muscular activity related to the production of sequences of gestures (Scott, 2008) could be a general limit on energetic investment. As a result, Menzerath’s law appears to emerge across communicative contexts.

There are a number of potential reasons for why language laws appear variable in their expression within ape gesture. For example, we might be considering the wrong unit of analysis. In human speech, sign, and gesture – as in other communication systems – it is possible to consider the production of a ‘unit’ of communication at different levels. For example, while Zipf’s law is clearly expressed in the duration of male rock hyrax vocalisations, that is not the case for female vocalisations where Zipf’s law of brevity emerges when analysing call amplitude rather than duration (Demartsev et al., 2019). Conversely, in Börstell et al. (2016) research on Swedish Sign Language, Zipf’s law of brevity seems to hold across sign categorisation, fingerspelling, and compounding. Interestingly, this study excluded the hold phase of a sign, limiting their analysis only to the more active stroke phase. The production of intentional gestures in apes are shaped not only by the signaller, but by the interaction between signaller and recipient (Byrne et al., 2017; Graham et al., 2022). As a result, the duration of hold or repetition phase may be shaped by the immediate context of the specific interaction – for example, in waiting for a response by the recipient it may vary between being absent and very prolonged. In contrast, the *action stroke* of a sign or gesture is always present and represents the need to convey information in that gesture, i.e., to discriminate it from other gesture actions. In Swedish Sign Language a prolonged and repetitive feedback sign and prolonged turn taking signs were the only two cases that diverged from the general Zipf’s pattern, as they were both long in duration as well as being highly frequent (Börstell et al., 2016). Zipf’s law acts on a signal ‘type’ in an individual’s or species’ repertoire – and it may be of interest to compare its expression across areas of gesture production that are more consistently produced across usage, such as the action stroke.

Research to date has typically focused on signal compression at the level of the communication system, but communication happens *in-situ*. Signallers likely respond to pressures on signalling efficiency more broadly: an intense but time-limited investment in clear signalling may be more energetically efficient than the need to travel with a female for extended periods following a failed signal. A similar solicitation with a different audience may need to be produced rapidly and inconspicuously, as the detection of this activity by other males could be fatal (Fawcett & Muhumuza, 2000). In a recent human study, pressures towards efficiency and accuracy were both required for Zipf’s law of brevity to emerge in experimental communicative tasks between two participants (Kanwal et al., 2017). Conversely, when participants were required to produce solely time-efficient vs solely accurate communicative signals no pattern emerged. The sexual solicitation context tested in our study may mirror the pattern seen in the time-efficient paradigm in the human study. In play, where urgency and time-efficiency may be less relevant, the same signals used by the same chimpanzees did show compression. While many vocalizations are relatively fixed (Janik & Slater, 1997; Fitch et al., 2016), gestural flexibility (in goal and context – Bard et al., 2019; Call & Tomasello, 2007; Hobaiter & Byrne, 2011a; Liebal et al., 2004) allows us to explore how compression acts within both specific instances of communication as well as on whole communication systems. To do so will require large longitudinal datasets in which it is possible to test both between-individual variation and within-individual variation across different gesture types and sequence lengths. Similarly, there remains substantial work needed to explore variation across different socio-ecological contexts of gesture use, for example in the social relationship between the signaller and recipient (Graham et al., 2022). The use of redundancy within specific subsets of gestural repertoire, or within specific contexts of gesture demonstrates both the importance of compression in communicative systems in general, but also the flexibility present in each specific usage. In doing so, it highlights the importance of exploring the impact of individual and socio-ecological factors within wider patterns of compression in biological systems in evolutionary salient scenarios.

## Methods

We measured *N*=560 male to female sexual solicitation gestures from 173 videos recorded within a long-term study of chimpanzee gestural communication depicting 16 wild, habituated East African chimpanzees (*Pan troglodytes schweinfurthii*) from the Sonso community of the Budongo Forest Reserve in Uganda (1°35’ and 1° 55’N and 31° 08’ and 31°42’ E), collected between December 2007 and February 2014. Observations were made between 7.30am and 4.30pm with recording of gestures following a focal behaviour sampling approach (Altmann, 1974). Here, all social interactions were judged to have the potential for gesture, in practice any situation in which two chimpanzees were in proximity and not involved in solitary activities, were targeted. Where several potential opportunities to record co-occurred, preference was given to individuals from whom fewer data had been collected (with a running record of data collection maintained to facilitate these decisions).

During October 2007 to August 2009 a Sony Handycam (DCR-HC-55) was used. Here video was recorded on MiniDV tape. The challenges of filming wild chimpanzees in a visually dense rainforest environment meant that, at times, the start of gestural sequences was not captured on video. Where this occurred, it was dictated onto the end of the video and these sequences were not included in analysis. Similarly, sequences in which part of the sequence was obscured, for example where a chimpanzee moves through dense undergrowth, were also discarded. After 2009 video data were collected using Panasonic camcorders (V770, HC-VXF1) were used which have a 3-second pre-record feature that improves the ability to capture the onset of behaviour; however, the same procedure was used and any sequences where the onset of gesturing was not clearly captured continued to be discarded.

### Sexual solicitation gestures

Sexual solicitation gestures were defined as those gestures given by a male towards a female with the goal of achieving sex, usually accompanied by the male having an erection and the female being in oestrus (Hobaiter & Byrne, 2011a, 2012). We included solicitations in the context of sexual consortship; here a male gestures in order to escort a female away from the group to maintain exclusive sexual access, which can occur prior to the peak of the female oestrus (Tutin, 1979). We restricted our analyses to male to female sexual solicitation, as female to male sexual solicitation attempts rarely involved sequences of gestures in this population. We further restricted analysis to solicitations by male individuals of at least 8-years old, as this is the minimum age of siring recorded in this community, limiting our signals to those on which there is more direct selective pressure.

### Defining gesture types and tokens

In quantitative linguistics, word *types* are used to assess Zipf’s law of brevity, whereas *tokens* are used to assess patterns conforming to Menzerath’s law. To distinguish the two, consider the question:

#### Which witch was which?

The question is composed of 4 *tokens* (overall word count), and three different word *types*, (which, witch, was). Gesture *types* (see S4 Table for a detailed repertoire description) were categorized according to the similarity of the gesture movement, which could be used either as a single instance or in a sequence; and each gestural instance represented an individual *token*.

Great apes deploy gestural sequences in two distinct forms (Hobaiter & Byrne, 2011b): one is the addition of further gestures following response waiting and is typically described as persistence (which may include elaboration). The second is the production of gestures in a ‘rapid sequence’ – here gestures are produced with less than 1 second between consecutive gesture tokens, and do not meet behavioural criteria for response-waiting occurring within a sequence (although it may occur at the end of it). As the expression of Menzerath’s law is typically considered at the level of a unique sequence, rather than one generated through the addition of gestures in response to earlier failure, we limit our analyses here to rapid sequences only. Sequence length was quantified as the number of gesture tokens produced with less than 1s between two consecutive gesture tokens; single gestures were coded as sequences of length one (Heesen et al., 2019; Hobaiter & Byrne, 2011b).

### Gesture duration

Gesture duration was calculated using MPEG streamclip (version 1.9.3beta). We measured gesture duration in frames, each lasting 0.04s. Gestural ‘units’ – like many other signals – can be considered at different levels of analysis, for example: a word is composed of syllables, and syllables of phonemes. Gestures have been described as composed of a preparation, action stroke, hold or repetition, and recovery phase (Kendon, 2004). Here we follow previous work in (Heesen et al., 2019) in defining the start of a gesture token as the initial movement of a part of the body required to produce the gesture. The end of a gesture token corresponded to (1) the cessation of the body movement related to gesture production, or (2) a change in body positioning if the gesture relied on body alignment, or (3) the point at which the goal was fulfilled, and any further movement represented effective action (for example, locomotion or copulation). Where the expression of a gesture token did not include a full recovery (in which the body part involved is returned to a resting state), the end of a token was discriminated from subsequent tokens through (1) a change in gesture action, e.g., from a reach to a shake, (2) a change in the rhythm or orientation of a gesture action, hold, or repetition, e.g., the rhythm or direction of an object shake is broken or changed (Hobaiter & Byrne, 2017).

### Intra-observer reliability

Video-based coding offers the opportunity to conduct reliability measures. Intra-observer reliability was tested by randomizing the order of the videos and re-coding the duration of the gestures of every ninth clip, for a total of 75 gestures from 23 clips. We performed an intraclass correlation coefficient (ICC) test – class 3 with *n*=1 rater (Landers, 2015) – which revealed very high agreement on gesture duration measurements (ICC=0.995, *p*<.001). Unfortunately, an additional step of inter-observer reliability was not possible due to the loss of the file that linked the original dataset to the videos from which data were extracted.

### Statistical analysis

All data were analysed using R version 4.0.0 and RStudio version 1.2.5042 (R Core Team, 2020; RStudio Team, 2020).We fitted Bayesian generalised linear multivariate multilevel models using the ‘brm’ function from the ‘brms’ package (Bürkner, 2017) with minimally informative priors, 2000 iterations and 3 chains.

We ran a first model testing Zipf’s law of brevity (Zipf-model), containing gesture token duration (s) as the response variable, the proportion of occurrences of a particular gesture type in the dataset (Proportion) as a fixed effect, and gesture Category (manual vs whole-body) as a control. We included signaller ID, sequence ID, and gesture type as random effects. We include Category as a variable here to allow for more direct comparison with previous work, which often excludes or differentiates non-manual signals, either in great ape gesture (Heesen et al., 2019; Rodrigues et al., 2021) or in signed languages and fingerspelling (e.g., Börstell et al., 2016).

We tested Menzerath’s law by running a second model (Menzerath-model) containing gesture token duration (s) as the response variable, sequence size (number of gesture tokens within the sequence) as a fixed factor, and the proportion of whole-body gestures within the sequence (PWB) as a control. We modelled signaller ID and sequence ID as random factors.

It was highlighted during the review process that the emergence of Menzerath’s law may be an artifice created by the selection of *fixed*, as opposed to *loose*, duration gesture types when producing longer sequences. To address this hypothesis, we produced histograms depicting the distribution of *loose* and *fixed* duration gestures within sequences at each sequence size. The majority of gesture types (n=20 of total 26), and of gesture tokens (n=456 of total 560) were of the *loose* gesture form, thus there were very few gesture sequences formed only of *fixed* gesture types. However, we further visually assessed the distributions of *fixed* gestures in sequences formed of only *fixed* gestures.

As our data may be particularly influenced by a single prolific individual (Duane) who contributed around half of the data, we assess the generalizability of our findings by replicating analyses conducted on the full dataset on a subset of the data containing only gestures by Duane as well as on a subset containing all but the prolific individual Duane. For the models testing Duane’s data, signaller ID was removed from the random factors as it was no longer relevant (with the inclusion of only one individual). In order to avoid inflation of the dataset we include date as a random factor; which also allows us to avoid biasing the analysis towards particularly prolific days and control for within-individual consistency.

We ran full-null model comparisons using the Level One Out information criterion (LOO) (Vehtari et al., 2017) ‘loo_compare’ function from the ‘stan’ package (version 2.21.5; Stan Development Team, 2022) where Zipf’s null model contained only the control variable Category and the random effects, whereas Menzerath’s null model contained only the control variable PWB and the random effects. Prior to the Bayesian analysis we assessed data distribution using the ‘fitdistr’ package (version 1.0-14; Delignette-Muller & Dutang, 2015). Following data inspection, we log-transformed gesture duration and average sequence duration as data from the response variable strongly skewed towards zero (for data inspection see supporting information 2).

Finally, previous work has frequently employed correlation and compression tests, which looks at whether the expected mean code length observed in the dataset is significantly smaller than a range of mean code lengths calculated via permutations, to test the mathematical theory behind both laws. In addition, we also fitted Bayesian generalised linear multivariate multilevel models with same number of iterations and chains as the previous models but having the median duration of each of the 26 gesture types as response variable, category of gesture as a fixed factor, as well as frequency of that gesture type as a predictor. These tests offer limited opportunities to control for potential confounds such as signaller identity and should be interpreted with caution in relatively small and variable datasets. We provide them in the supporting information 6 to allow for comparison with previous work that analysed median durations with or without implementing generalised linear models (e.g., Hernández-Fernández et al. 2019; Watson et al. 2020).

### Data and code

Data and code for all analyses are available in a public GitHub repository: github.com/Wild-Minds/LinguisticLaws_Papers

## Acknowledgements

We thank the staff and field assistants of the Budongo Conservation Field Station for their assistance in the original gestural data collection, and the Ugandan National Council for Science and Technology and the Ugandan Wildlife Authority for permission to conduct the original research. We thank the Royal Zoological Society of Scotland for its funding of the field station. We thank Dr Alexander Mielke for his advice on the statistical models. We thank the editor and three anonymous reviewers for their constructive comments. This research received funding from the European Union’s 8^th^ Framework Programme, Horizon 2020, under grant agreement no 802719.

## Supporting information 1 – Gesture types definitions

**Table S1.**
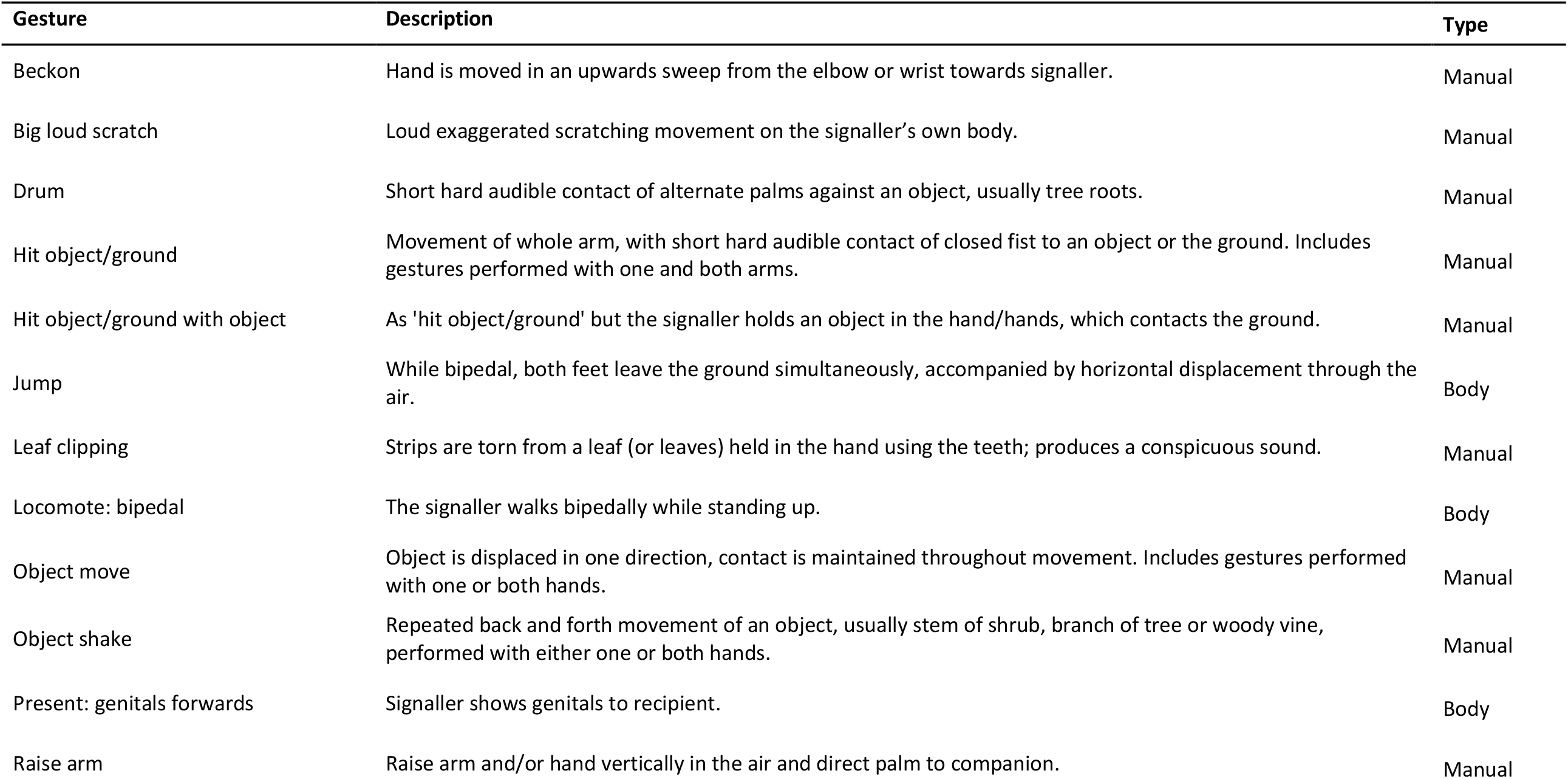
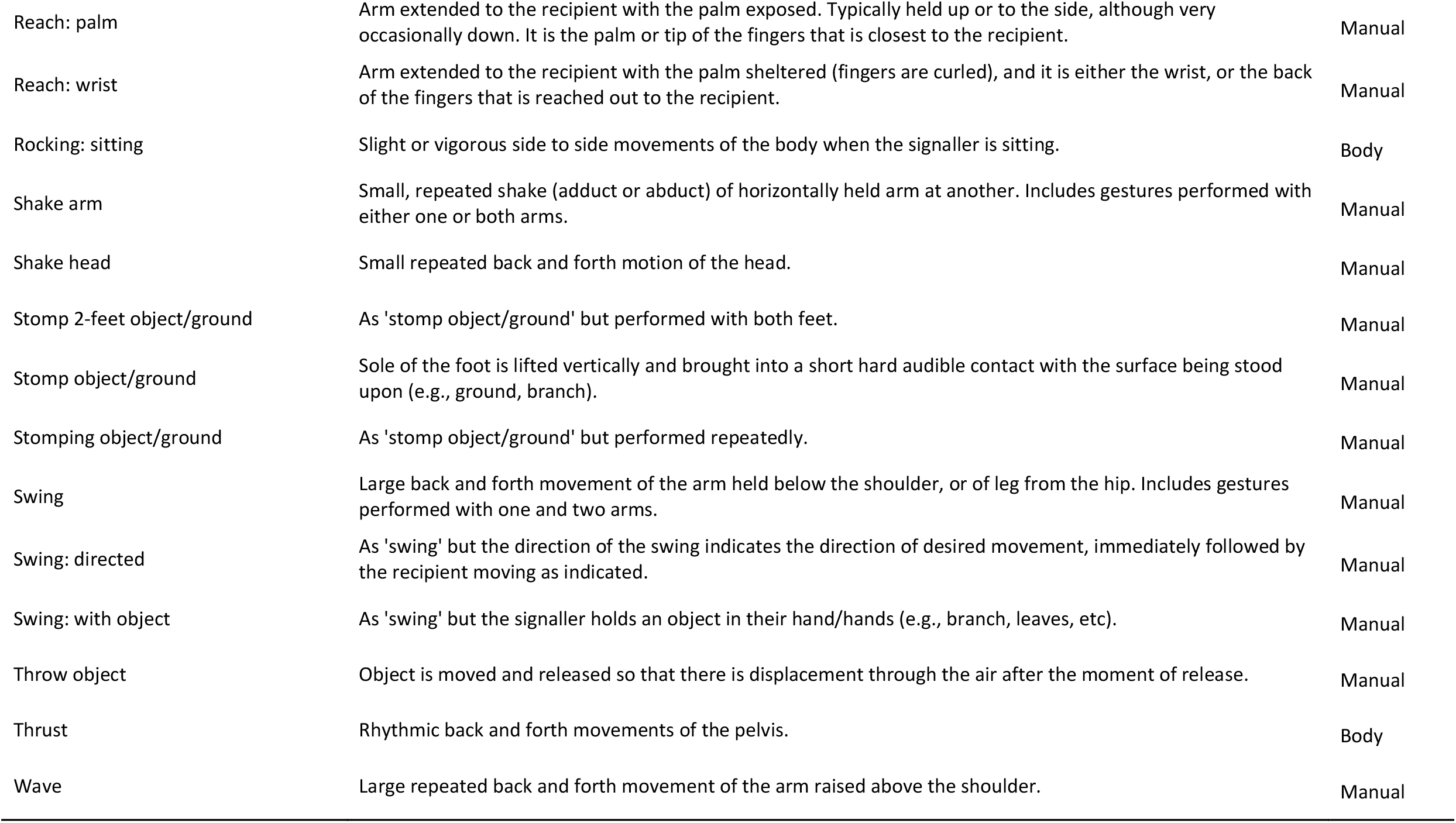
Ethogram of the 26 gesture types recorded in the dataset. Definitions are taken from (Hobaiter & Byrne, 2011a) and (Nishida, 2010). Video examples and illustrations of these gestures are available at www.greatapedictionary.com.

## Supporting information 2 - Duration distribution analysis

Before performing the GLMM analysis we analysed the distribution of the gesture duration data by (1) visually inspecting its empirical density and cumulative distribution (Figure S4) and (2) assessing its skewness and kurtosis via the visual inspection of the Cullen and Frey graph (Figure S5). Figure S5 shows the data are skewed towards low values, as almost half of the data lays between 0 and 3 seconds. Further, we fitted three theoretical distributions to the data – namely Weibull, Gamma, and Lognormal – and compared loglikelihood values (Table S2). We then plotted the three distributions and visually inspected the Q-Q, P-P, and histogram density plots (Figure S6). Finally, we compared Weibull, Gamma, and Lognormal distributions against gesture duration data distribution via goodness-of-fit tests and goodness-of-fit information criterion (Table S3 and S4), which helped identify the lognormal distribution as the best fitting one. Therefore, we proceeded with log-transforming the duration variable to best fit model assumptions.

**Figure S1.**
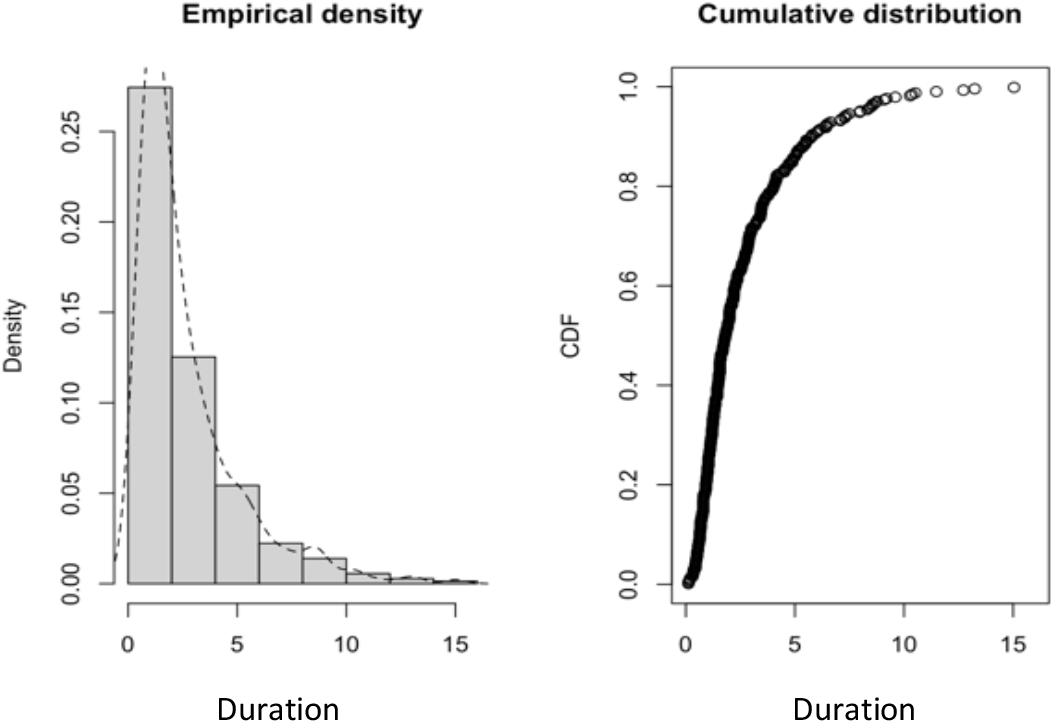
Empirical distribution of gesture duration. Histogram and empirical cumulative distribution function (CDF) plots representing the distribution of gesture duration. Histogram bars represent sample distribution, dashed line indicates empirical density.

**Figure S2.**
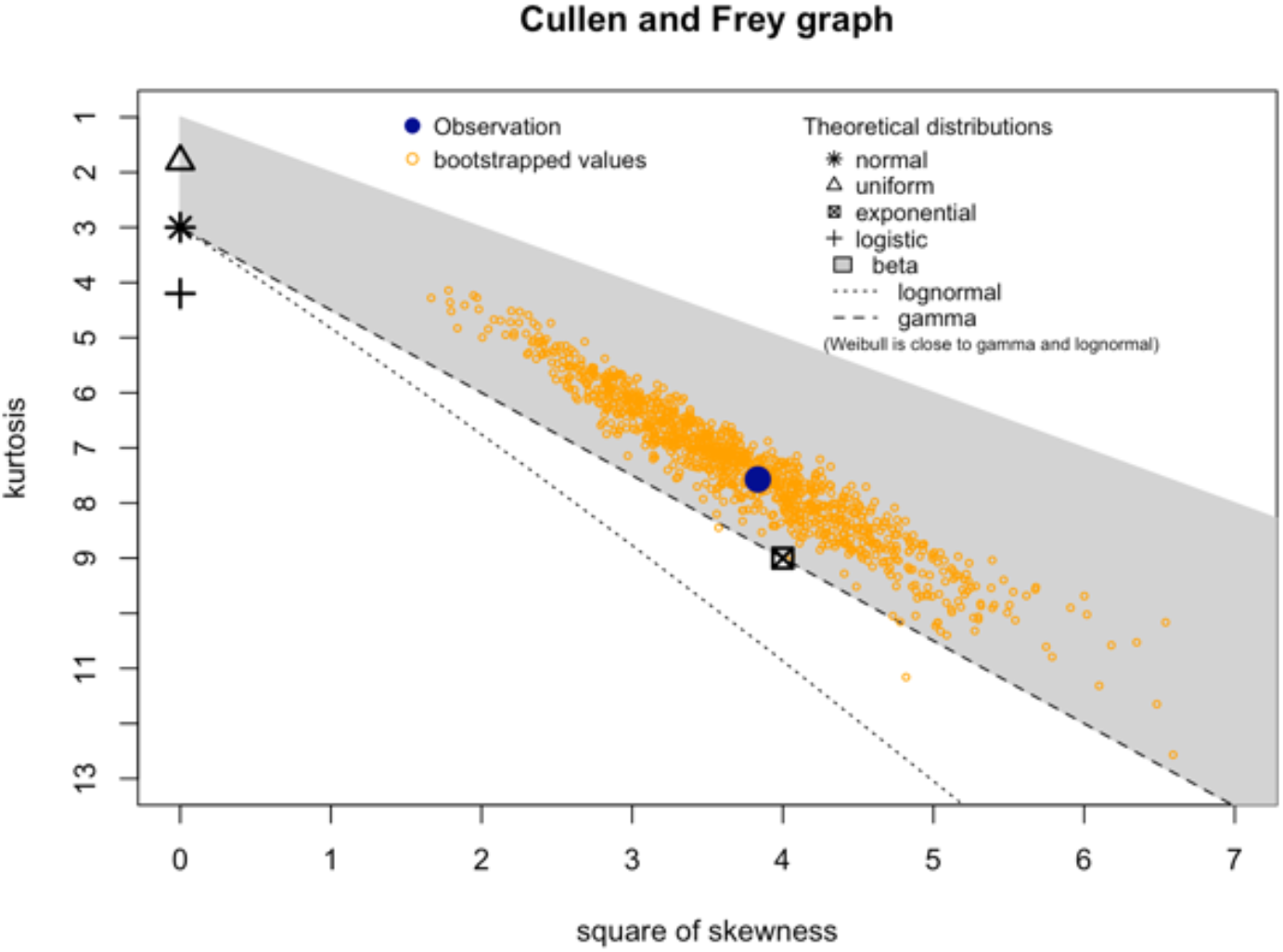
Cullen and Frey graph for gesture duration. The graph depicts the distribution of the skewness and kurtosis of gesture duration data with bootstrapped values, plotted against other theoretical distributions, namely normal, uniform, exponential, logistic, beta, lognormal, and gamma.

**Table S2.**
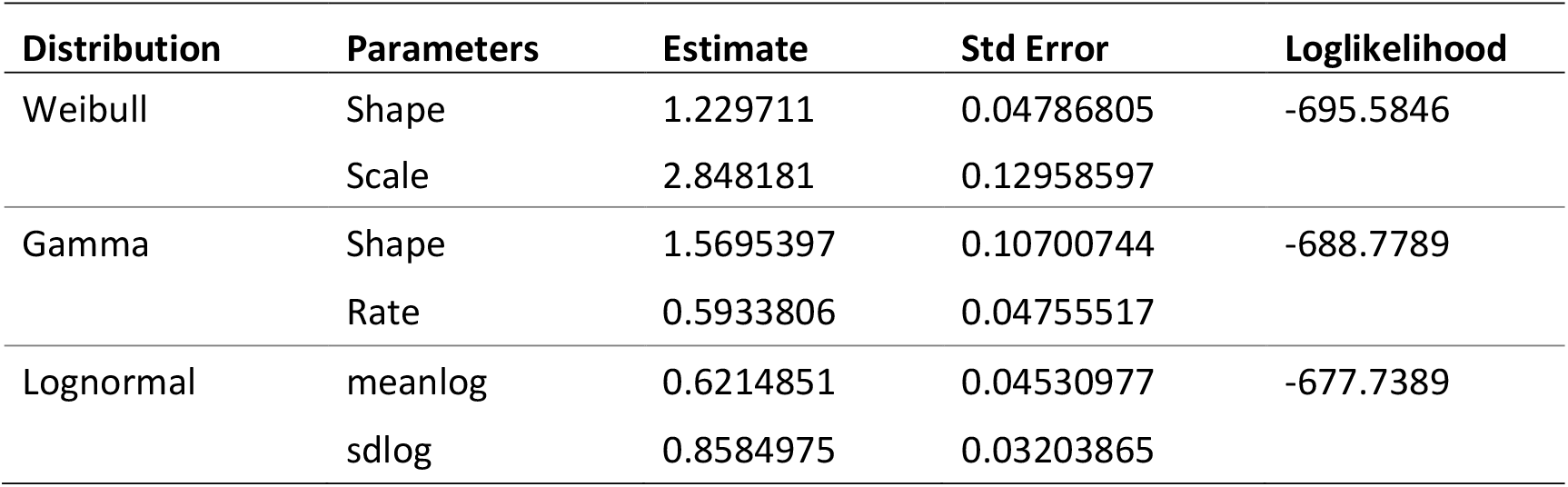
Estimate and standard error for fitting the parameters of three theoretical distributions to the distribution of the gesture duration data.

**Figure S3.**
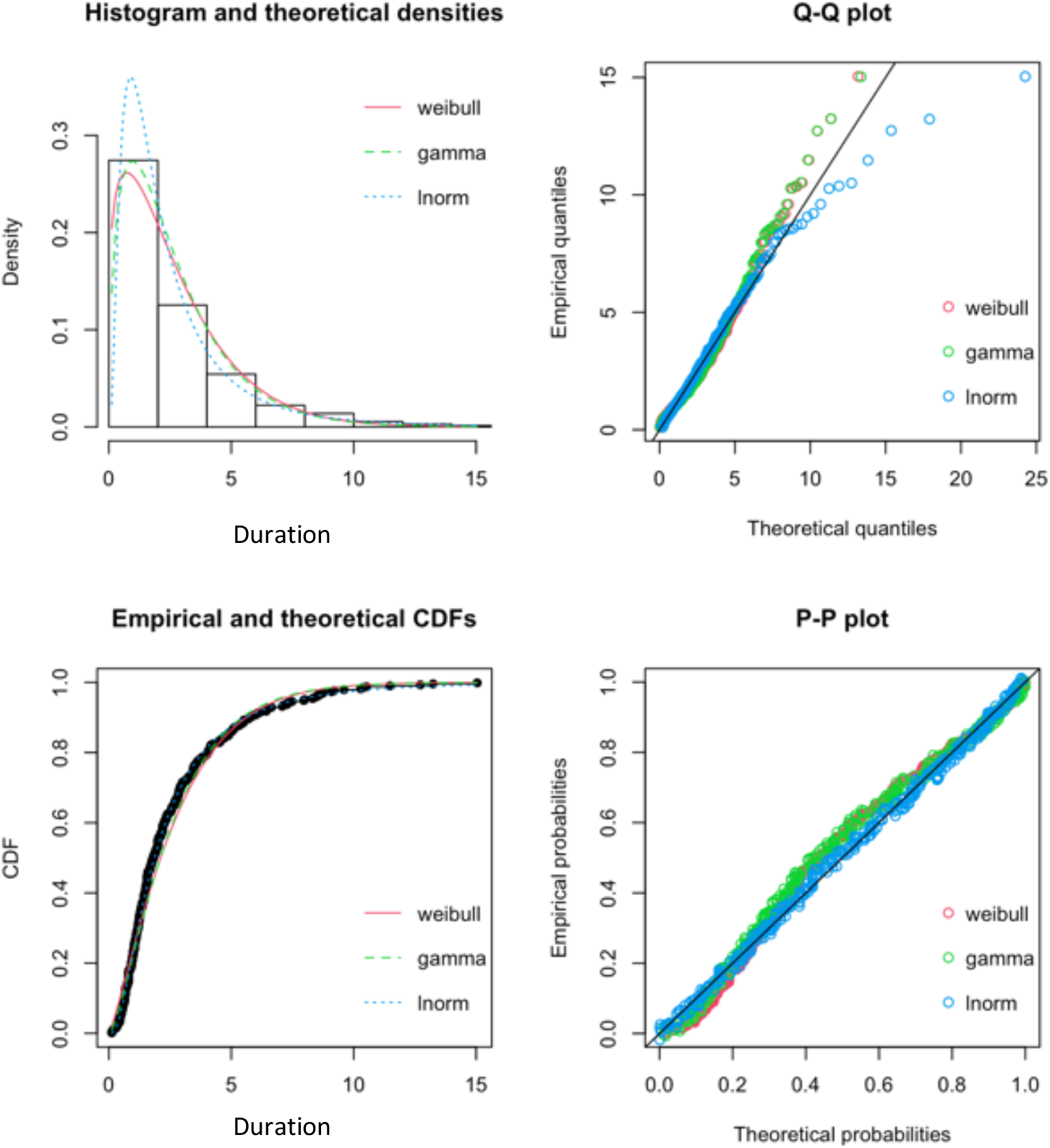
Histogram and theoretical densities, Q-Q and P-P plots depicting the gesture duration data distribution against the fitted Weibull, Gamma, and Lognormal distributions. Histogram represents the distribution of duration data while the red, dashed green, and dashed blue lines indicate the theoretical Weibull, Gamma, and Lognormal distributions, respectively.

**Table S3.**
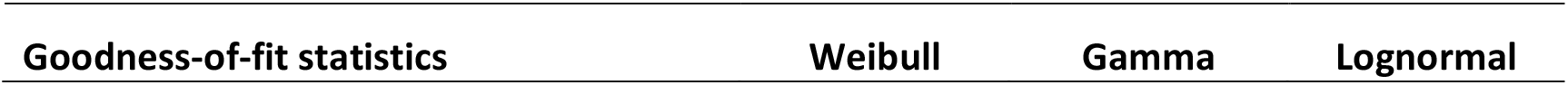
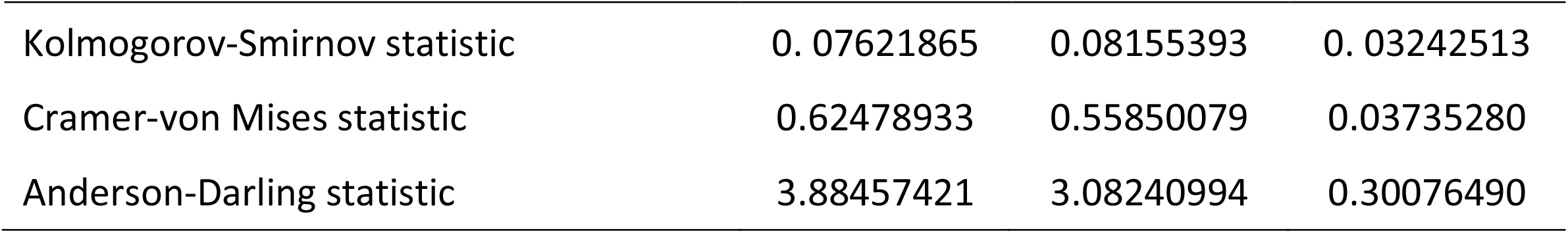
Goodness-of-fit statistics compared across fitted distributions to the gesture duration data.

**Table S4.**
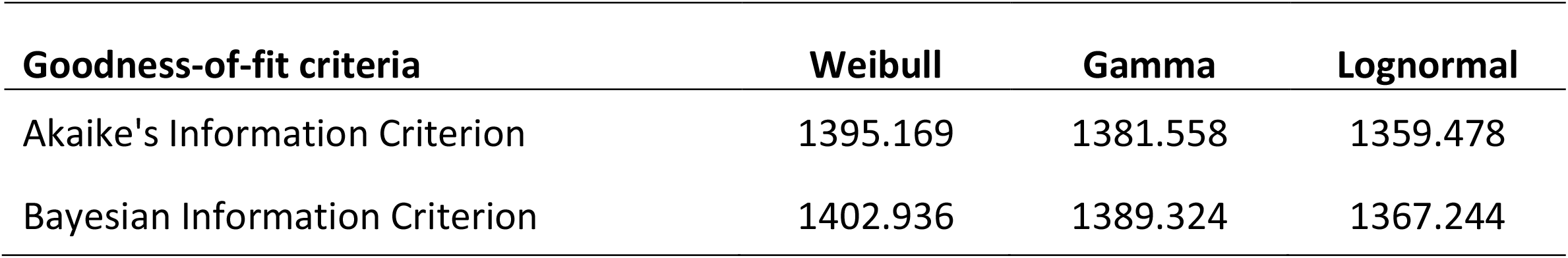
Goodness-of-fit information criteria compared across fitted distributions.

## Supporting information 3 - Model results

**Table S5.**
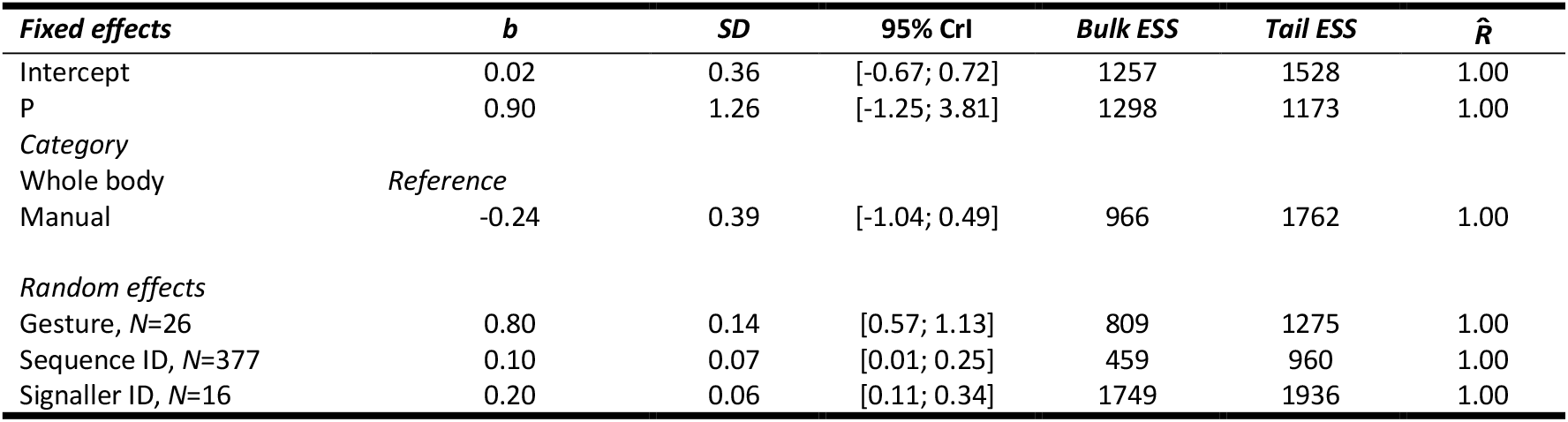
Summary of the Bayesian mixed model analysis results for the Zipf-model which included all the data (*N*=560).

**Table S6.**
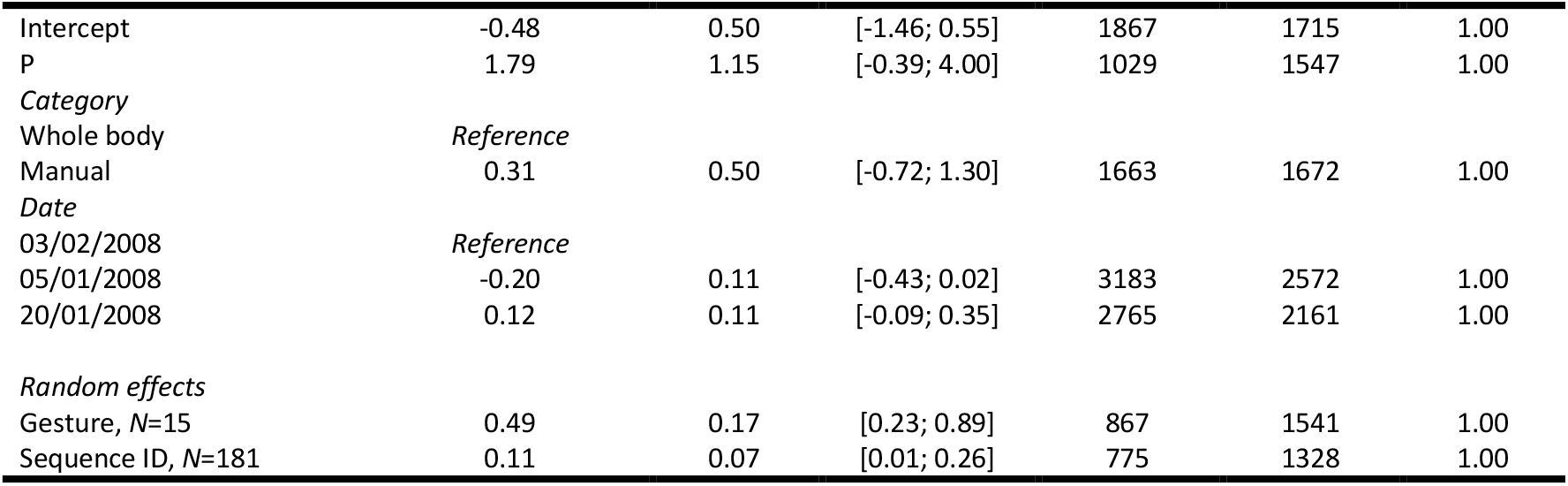
Summary of the Bayesian mixed model analysis results for the Zipf-model which included only Duane’s data (*N*=290).

**Table S7.**
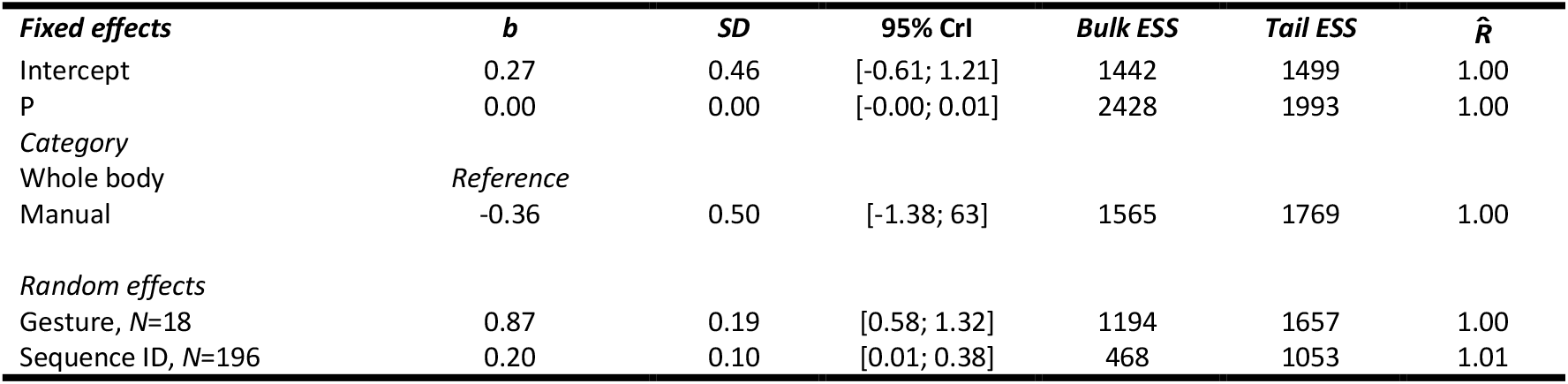
Summary of the Bayesian mixed model analysis results for the Zipf-model which included data from all individuals but Duane (*N*=270).

**Table S8.**
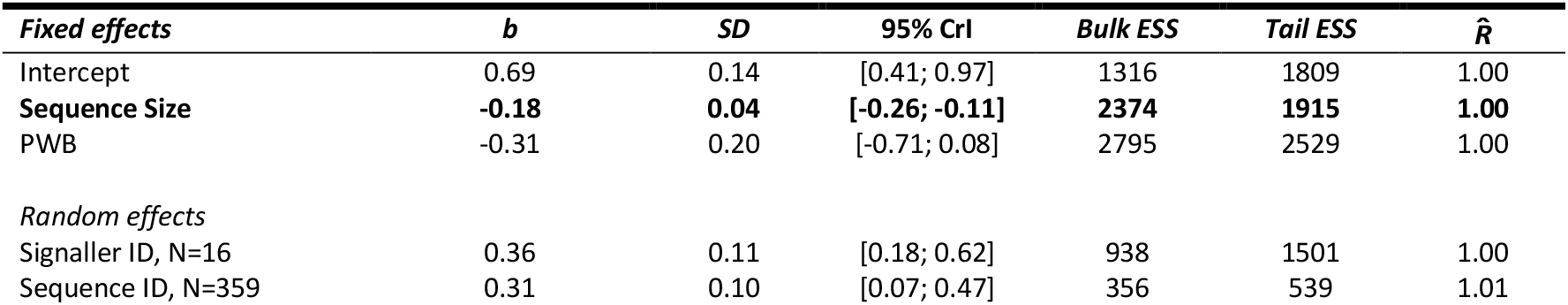
Summary of the Bayesian mixed model analysis results for the Menzerath-model which included all the data (*N*=530).

**Table S9.**
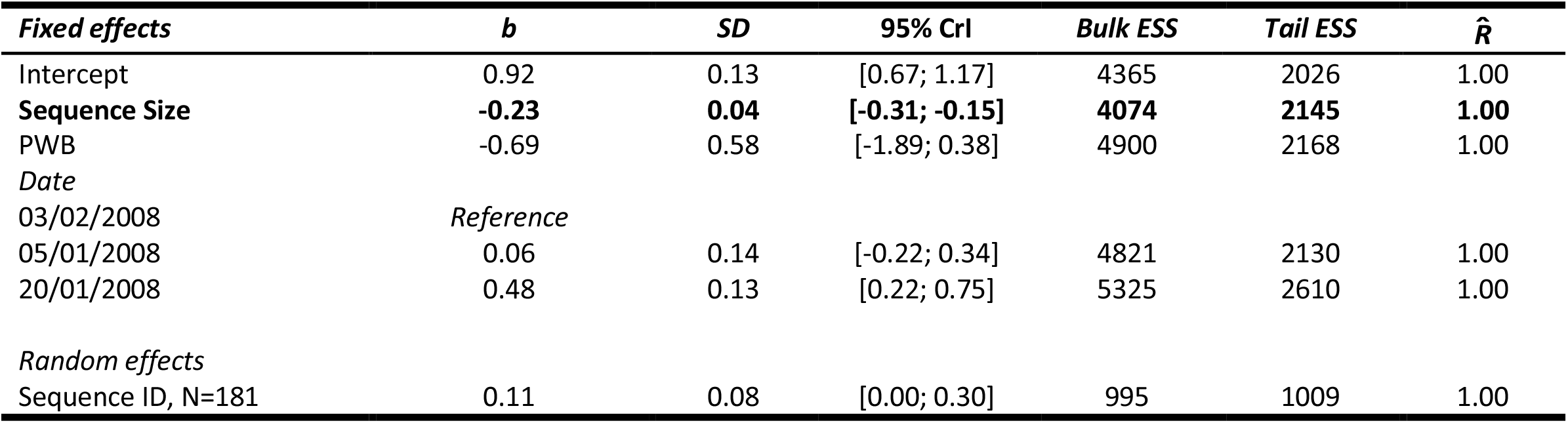
Summary of the Bayesian mixed model analysis results for the Menzerath-model which included only Duane’s data (*N*=273).

**Table S10.**
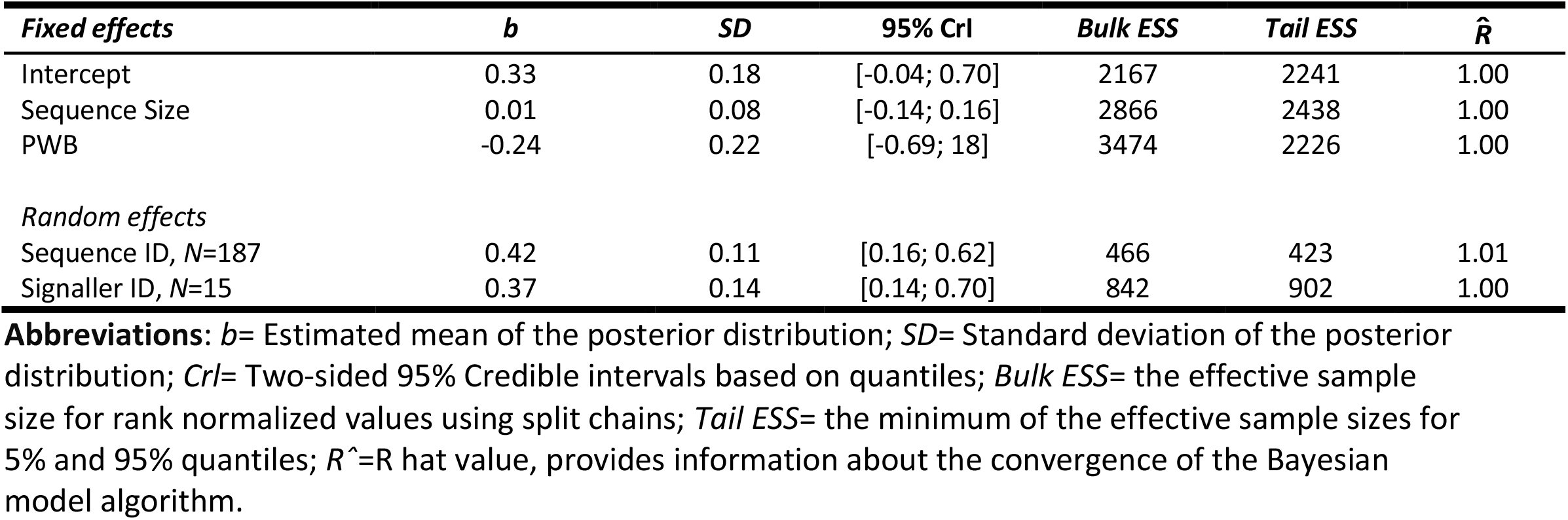
Summary of the Bayesian mixed model analysis results for the Menzerath-model which included data from all individuals but Duane (*N*=257).

## Supporting information 4

**Figure S4.**
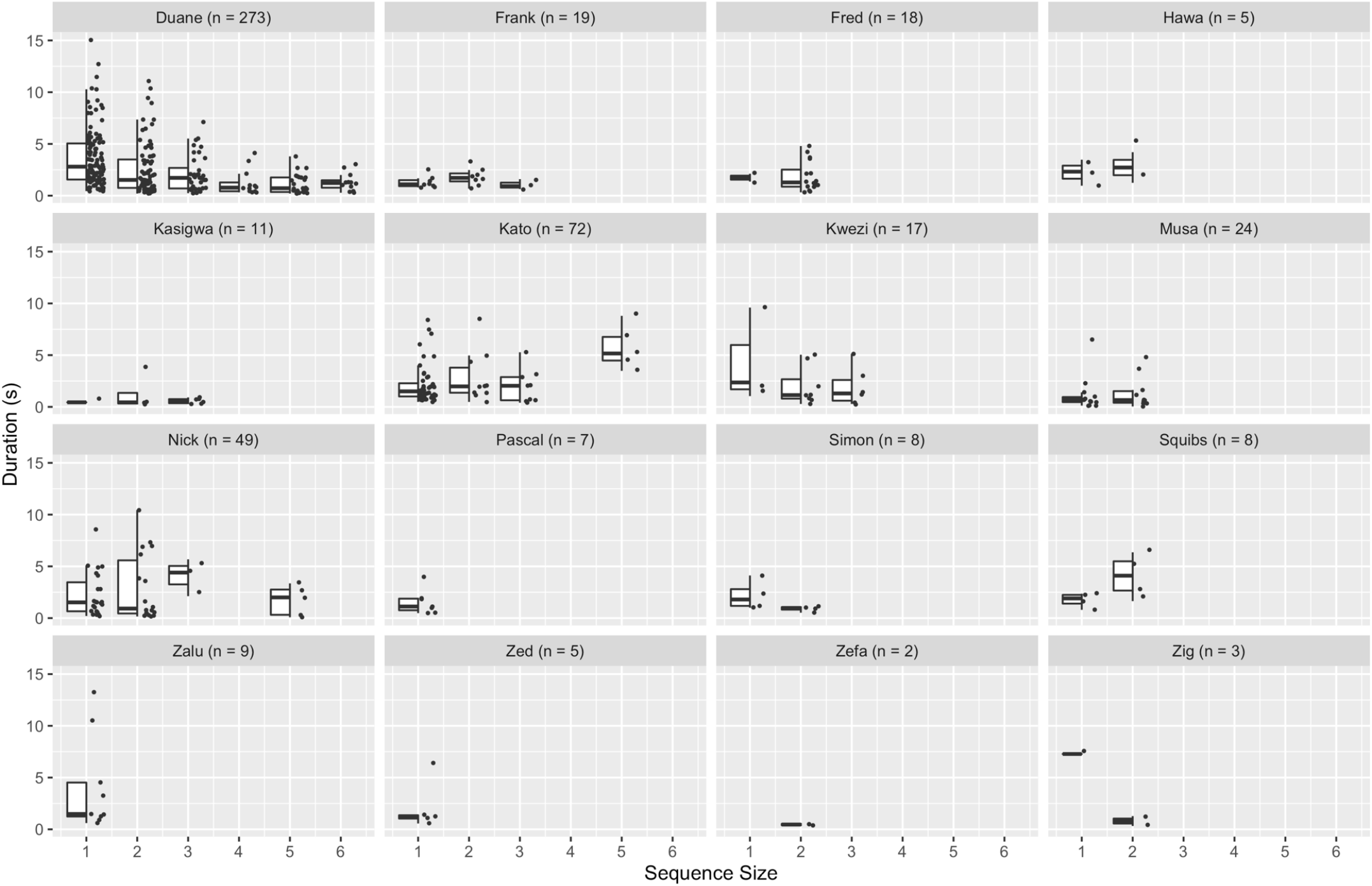
Distribution of gesture durations based on sequence size for each of the 16 individuals in the dataset. Points represent individual gesture tokens. Boxplots show median (black central bar), interquartile range (boxes), maximum and minimum values exploding outliers (whiskers). n indicates sample size for each individual.

## Supporting information 5 – Visual inspection of sequence structure

**Figure S5.**
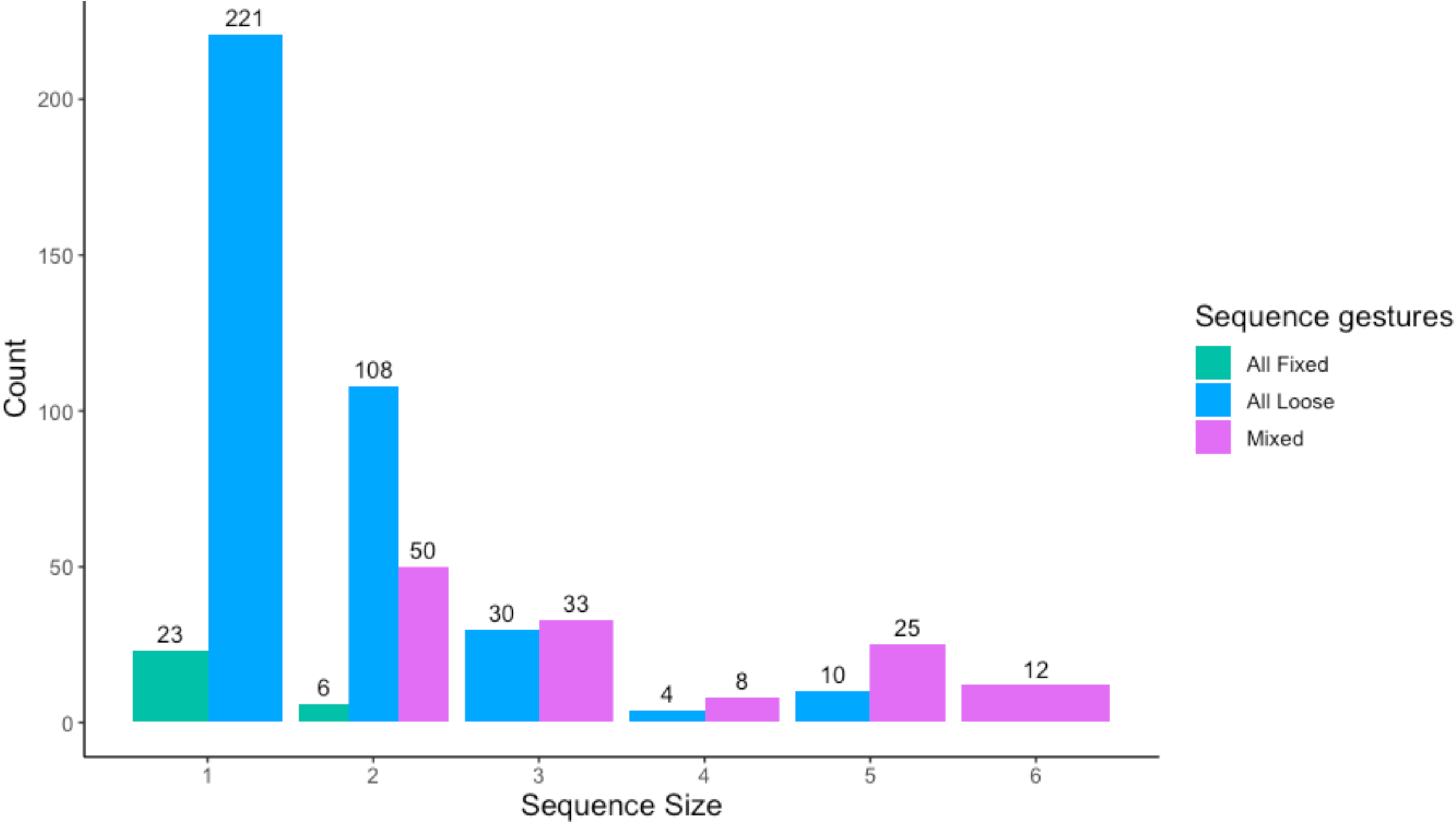
Bar chart showing the frequency distribution of the three different types of sequences depending on their sequence size. Green sequences comprise only *fixed* duration gestures, blue sequences only *loose* duration gestures. Pink bars represent sequences formed by a mix of *loose* and *fixed* duration gestures. Numbers above bars indicate the frequency of each sequence type per sequence size.

**Figure S6.**
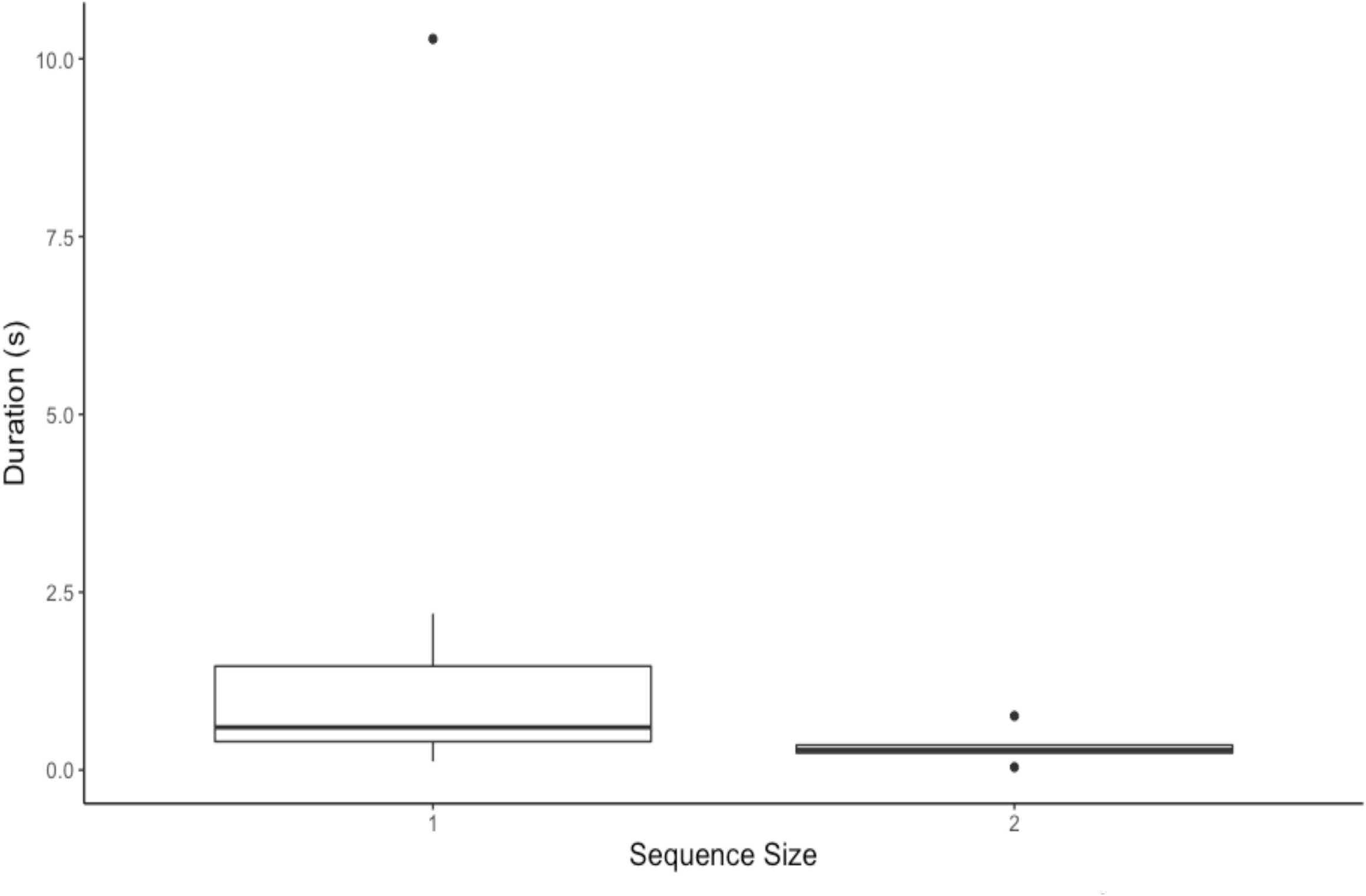
Boxplots of the duration of gestures with constrained duration (i.e., *fixed* duration gestures) in sequences formed of *fixed* duration gestures solely. Boxplots show median (black central bar), interquartile range (boxes), maximum and minimum values exploding outliers (whiskers) and outliers (circles).

## Supporting information 6 – Alternative analyses

### Correlation and compression

#### Methods

Compression predicts that mean duration should be smaller than expected by chance (Ferrer-i-Cancho et al., 2013). Similarly, optimal compression predicts linguistic laws as a correlation in a specific direction, *i*.*e*., the correlation cannot be positive (Ferrer-i-Cancho et al., 2013, 2020). Accordingly, we employed one-tailed tests of compression throughout, but we also report the outcome of two-tailed equivalents for comparison with previous findings (Heesen et al., 2019).

We conducted one-tailed Spearman rank correlation tests to analyse the relationship between the frequency within the sample of a gesture type (*frequency*) and its mean duration (*mean gesture type duration*), calculated by dividing the total sum of all durations of the same gesture type (Sum), by *frequency (i*.*e*., *duration=Sum/frequency)* (Semple et al., 2013). A similar procedure was used to test for a correlation between the mean gesture duration within a given sequence (*sequence*) and the number of gesture tokens in the same sequence (*n*). Mean gesture duration was calculated by dividing the total duration of a gestural sequence (*Total*) – *i*.*e*., the sum of all durations of the gesture tokens in the sequence excluding pauses between gestures – by the number of gesture tokens within that sequence *n* (*i*.*e*., *mean gesture duration within sequence*=*sum of durations of gestures within the sequence*/*number of gestures within the sequence*). A negative correlation between *mean gesture type duration* and *frequency* coherent with Zipf’s law of brevity, and a negative correlation between *t* and *n* conforming to Menzerath’s law could both be unavoidable artefacts given the relationship between *d* and *f*, and between *t* and *n –* as defining *d* involves *f*, and defining *t* involves *n* – which could lead to *d* = 1/*f* and *t*=1/*n* (Ferrer-i-Cancho et al., 2014). Such artefacts can be excluded by establishing that *D* and *f*, and *T* and *n* are significantly positively correlated (Ferrer-i-Cancho et al., 2014; Semple et al., 2013), which we tested using two Spearman rank correlation tests. Current findings suggest that the expressions of linguistic laws are not ‘universal’, and there may be more variation than previously recognised (Semple et al., 2022). For example: earlier research demonstrated Zipf’s law of brevity can be present in parts of a repertoire, when it appears to be absent in the whole repertoire (Ferrer-i-Cancho & Hernández-Fernández, 2013; Heesen et al., 2019). As a result, we also tested for Zipf’s law of brevity in specific subsets of the repertoire, namely manual versus whole-body gesture types as these had been found to differ in previous work (Heesen et al., 2019). Moreover, a specific check of Zipf’s law of brevity in manual gestures aids in comparison with studies of human communication that only consider manual signals (for example in signed languages and fingerspelling).

### Compression test

#### Is the mean duration of chimpanzee sexual solicitation gesture types significantly small?

Following earlier work on chimpanzee play gestures (Heesen et al., 2019), we first calculated mean duration of all gesture types *L* via the equation:

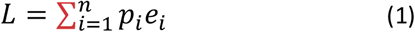

where *n* is the number of elements within the repertoire, *p*_*i*_ is the normalized probability of the *i*^*th*^ element – calculated by dividing the frequency of the *i*^*th*^ gesture by the total frequency of all gestures – and *e*_*i*_ is the magnitude of the *i*^*th*^ element (*i*.*e*., its average duration *d*).

To test for compression and whether Zipf’s law holds in chimpanzee sexual solicitation gestural communication, we used a permutation test assessing whether *L* was significantly small (Sokal & Rohlf, 1995). Following (Heesen et al., 2019) we created “*a control distribution of L (L’) defined by a permutation function π (i)”* and calculated the left *p*-value by dividing the number of permutations where *L’*≤*L* by the number of total permutations, here 10^5^. *L* was also calculated and tested for each subset created.

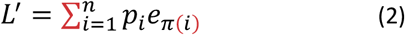

#### Is the expected total sum of the duration of gestures of each sequence significantly small?

As explained by (Heesen et al., 2019), the total duration of a collection of sequences can be quantified as

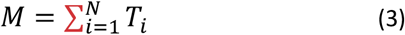

Where *T*_*i*_ is the total duration of the *i*th sequence and *N* is the number of sequences. In turn,

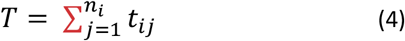

where *t*_*ij*_ is the duration of the *i*th element of the *i*th sequence and *n*_*i*_ is the size of the *i*th sequence. Given that the mean duration of gestures from the *i*th sequence can be expressed as 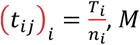, *M* can be defined as

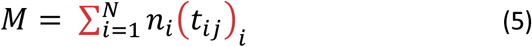

*M* was calculated through this equation and was tested to assess whether it is significantly small. We performed a similar permutation test to that conducted to test for the significance of *L*, to check whether *M* was significantly small as compared to the values generated by random permutation of the data (Zipf, 1936). In such case, *n*_*i*_ has the role of *p*_*i*_ and (*t*_*ij*_)_*i*_ has the role of *e*_*i*_ in the test, with *n*_*i*_ and (*t*_*ij*_)_*i*_ remaining constant during the test.

The permutation test produces a left *p*-value to check if *L* (or *M*) is significantly small and a right *p*-value to check if *L* (or *M*) is significantly large compared to the distribution of the values created by a permutation of the data (Heesen et al., 2019). The total number of permutations carried out was *R*=10^5^.

### Results of one-tailed analyses

#### Zipf’s law of brevity

##### Do chimpanzee sexual solicitation gestures follow Zipf’s law of brevity?

We did not find a pattern in agreement with Zipf’s law of brevity; there was no evidence for a significant negative correlation between mean gesture type duration (*d*) and frequency of use (*f*) (Spearman correlation: *r*_s_=0.30, *n*=26, *p*=0.066), in agreement with the Bayesian model analysis. Consistent with this result, the compression test revealed that the expected mean code length of gesture types *L* had a magnitude of 2.39s and was not significantly small (*p*_*left*_=0.951). Rather, *L* was significantly big (*p*_*right*_=0.05, Figure S7).

**Figure S7.**
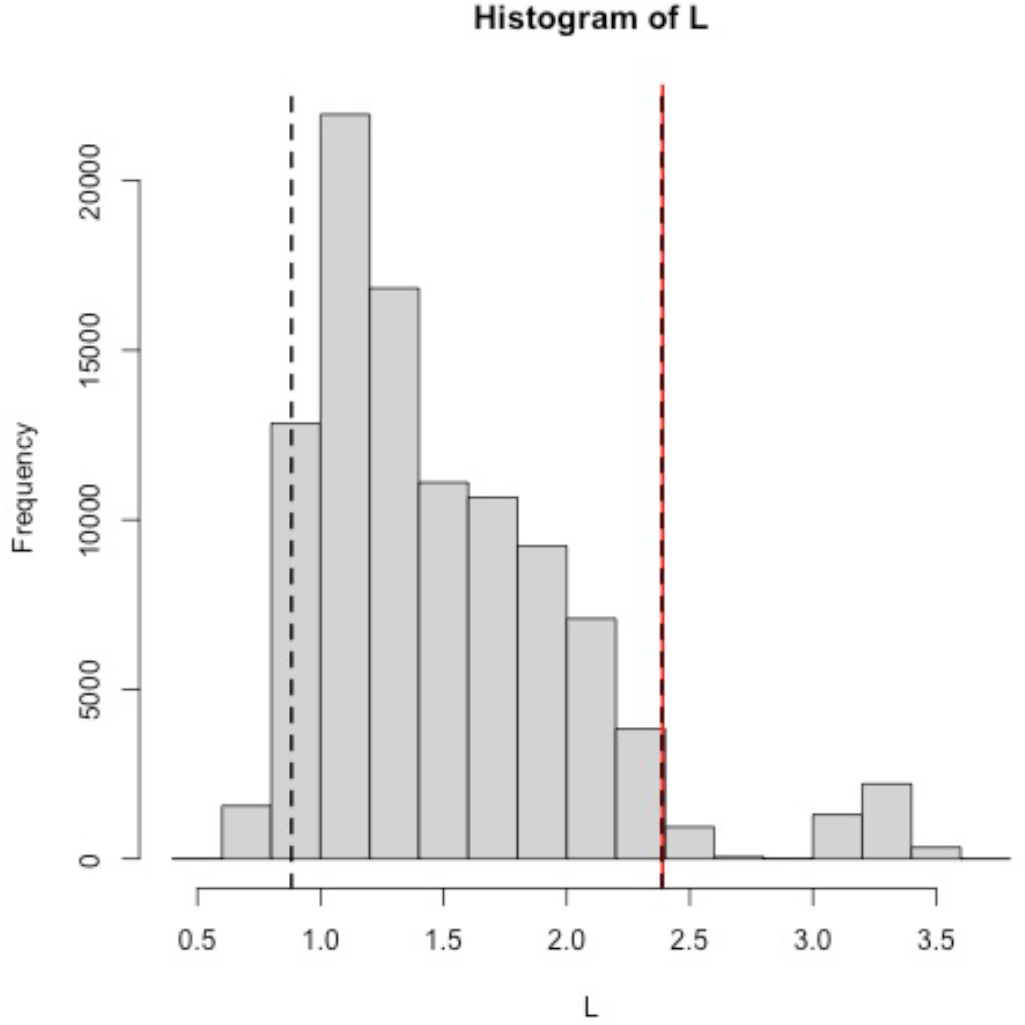
Histogram showing the distribution of the permuted *L* values. Observed *L* value is highlighted with the red continuous line. Black dashed line indicates the lower and upper 5% of the permuted data.

##### Subset analysis: whole-body and manual gesture types

We found no evidence for a negative correlation between *d* and *f* when separating whole-body gestures from manual gestures (Spearman’s rank correlation: whole-body, *r*_*s*_=-0.3, *n*=5, *p*=0.342; manual, *r*_*s*_ =0.42, *n*=21, *p*_*left*_=0.969). Rather, manual gestures showed a significant positive correlation (*r*_*s*_ =0.42, *n*=21, *p*_*right*_=0.031). Compression tests revealed that for whole-body gestures, *L*=0.13s and was neither significantly big or small (*p*_*left*_=0.174, *p*_*right*_=0.817), and for manual gestures, *L*=2.26s and, if anything, tended towards being significantly big (*p*_*right*_=0.058) rather than small (*p*_*left*_=0.942; Figure S8).

**Figure S8.**
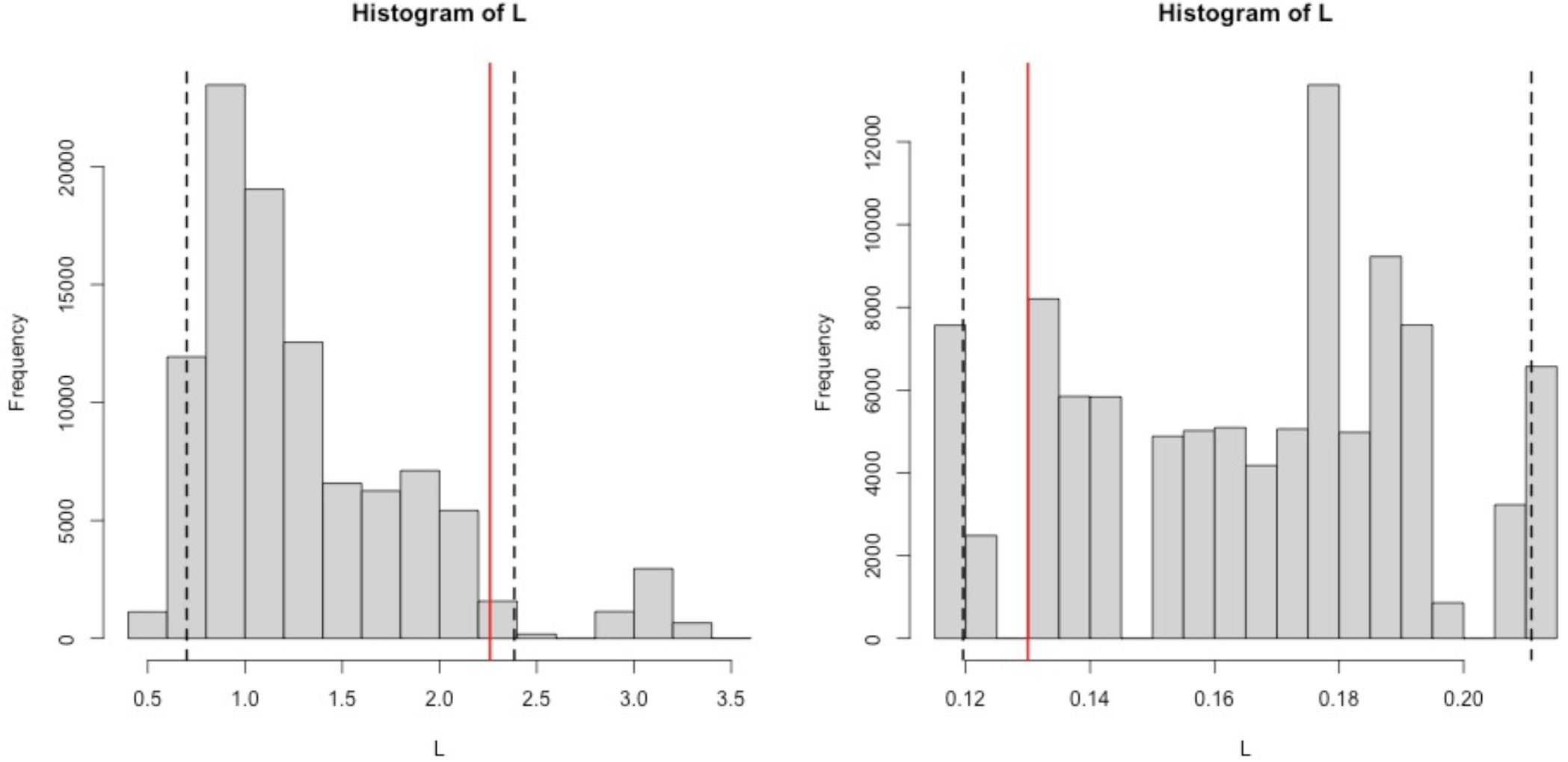
Histograms showing the distribution of the permuted *L* values for manual gestures (left) and whole-body gestures (right). Observed *L* value is highlighted with the red continuous line. Black dashed lines indicate the upper and lower 5% of the permuted data.

##### Do chimpanzee sexual solicitation gesture sequences follow Menzerath’s law?

We tested Menzerath’s law in 359 sequences, composed of 530 gesture tokens; there was no evidence for a negative relationship between mean constituent duration and sequence size (Spearman’s rank correlation: *r*_*s*_=-0.08 *n*=359, *p*=0.076). However, the compression test revealed that the total sum of the duration of each sequence *M* had a value of 1300.67 and was significantly small (*n*=359, *p*=0.003; Figure S9).

**Figure S9.**
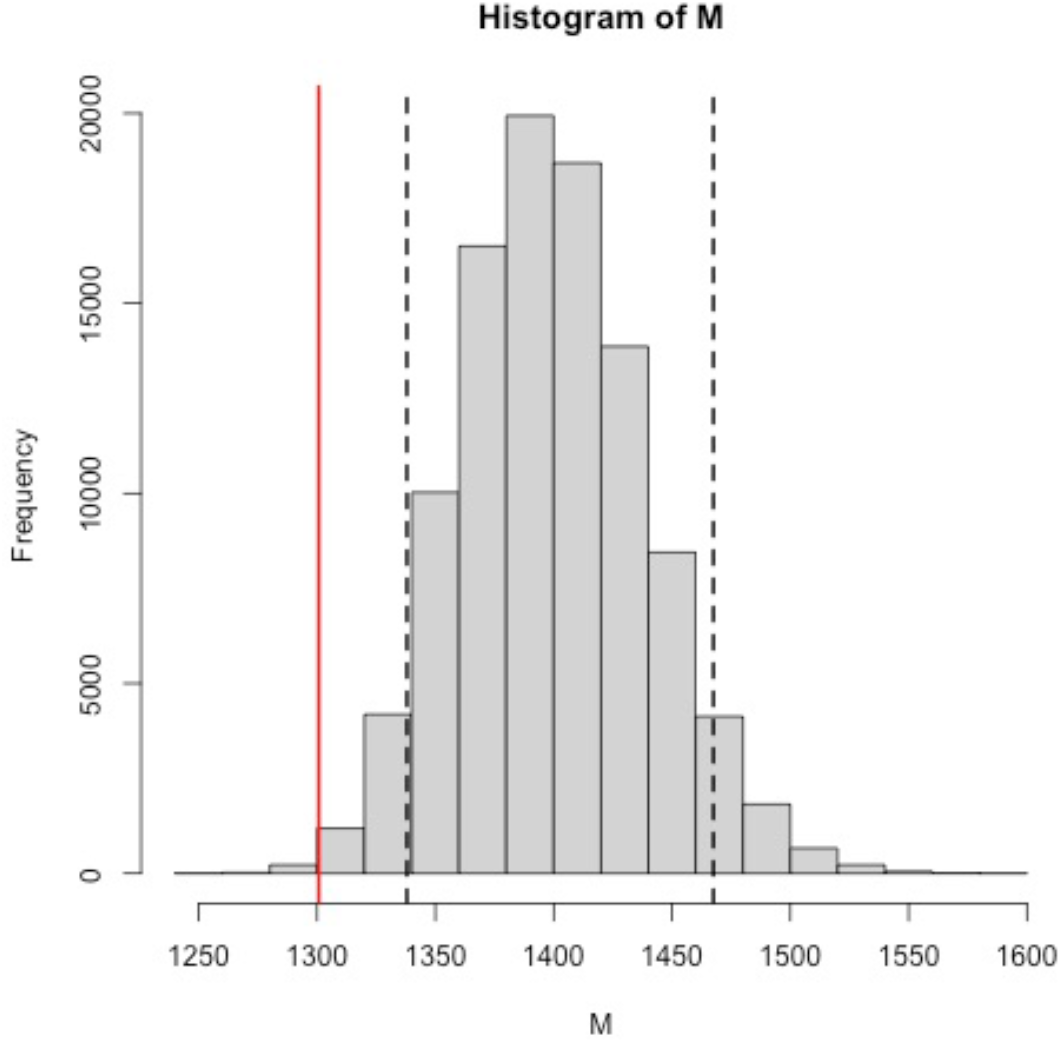
Histogram showing the distribution of the permuted *M* values. Observed *M* value is highlighted with the red continuous line. Black dashed line indicates lower 5% of the permuted data.

## Discussion

The results from the correlation analysis must be taken with caution as this analysis does not control for individual variation, gesture type, and sequence in which the gesture is performed. The Bayesian model that included these factors and which tested for Zipf’s law of brevity (Zipf-model) was similar to the respective null-model, suggesting that frequency of gesture type within the dataset and category of gesture type did not predict gesture duration.

The contrast between the correlation analysis and the compression analysis for Menzerath’s law highlight how individual variation may show an apparent absence of pattern in the correlation analysis but a strong effect in the Bayesian model analysis, where it is controlled for.

## Results of two-tailed analyses

### Zipf’s law of brevity

We did not find a pattern corresponding to Zipf’s law of brevity, with no correlation between mean gesture type duration (*d*) and frequency of use (*f*) (Spearman correlation: *r*_s_=0.30, *n*=26, *p*=0.131). When analysing only manual gestures, *f* and *d* tended to be significantly positively correlated (Spearman correlation: *r*_s_=0.42, *n*=21, *p*=0.061). Conversely, we did not find any correlation between *f* and *d* in whole body gestures (Spearman correlation: *r*_s_=-0.3, *n*=5, *p*=0.683).

### Menzerath’s law

We failed to find a pattern between sequence size *n* and mean constituent duration *t* of the same sequence that followed Menzerath’s law (Spearman correlation: *r*_*s*_=-0.08, *n*=376, *p*=0.142). When analysing sequences comprising only whole-body size and average gesture duration showed a significant positive correlation (Spearman correlation: *r*_*s*_=0.59, *n*=20 *p*=0.005). Sequence size and average gesture duration did not correlate in sequences composed of only manual gestures (Spearman correlation: *r*_*s*_=-0.06, *n*=315 *p*=0.324), or in those formed by both manual and body gestures (Spearman correlation: *r*_*s*_=0.09, *n*=24 *p*=0.673).

### Bayesian analysis on the medians of the 26 gesture types

We ran additional brms analysis computing the median duration per each gesture type across the whole dataset. Median duration of each gesture type was assigned as response variable, category of gesture as fixed factor as well as frequency of that gesture type as a predictor. We ran a model with 2000 iterations and 3 chains. The full model was no different from the null model which excluded the frequency of gesture type as a predictor (LOO difference: -0.3 ± 0.5; Table S11). We ran a similar analysis on the gestures performed by the one individual Duane, with similar results (full-null model comparison, LOO difference: -0.6± 1.2; Table S12). Please note that that individual identity is not controlled for in these analyses and they should be interpretted with caution.

**Table S11.**
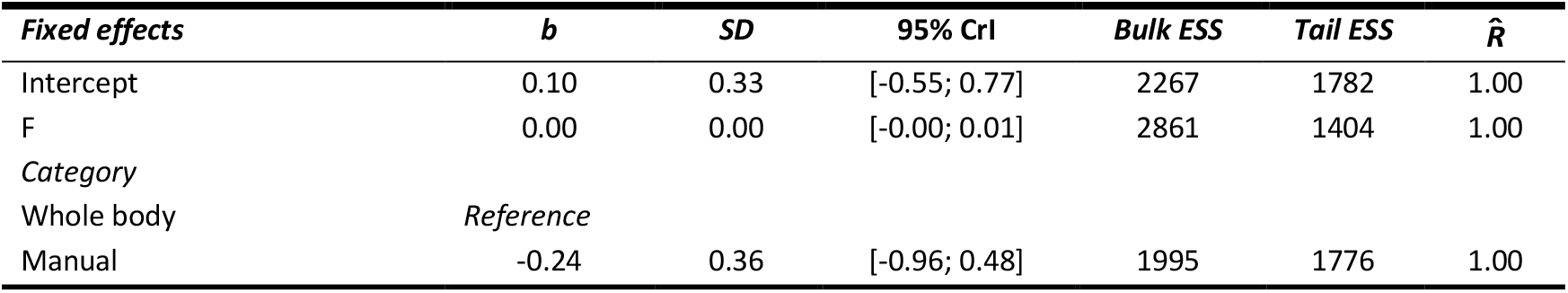
Summary of the Bayesian mixed model analysis results for the Zipf-model which included only a median value per gesture type as response variable, the frequency of gesture type as predictor and category of gesture type as control (*N*=26).

**Table S12.**
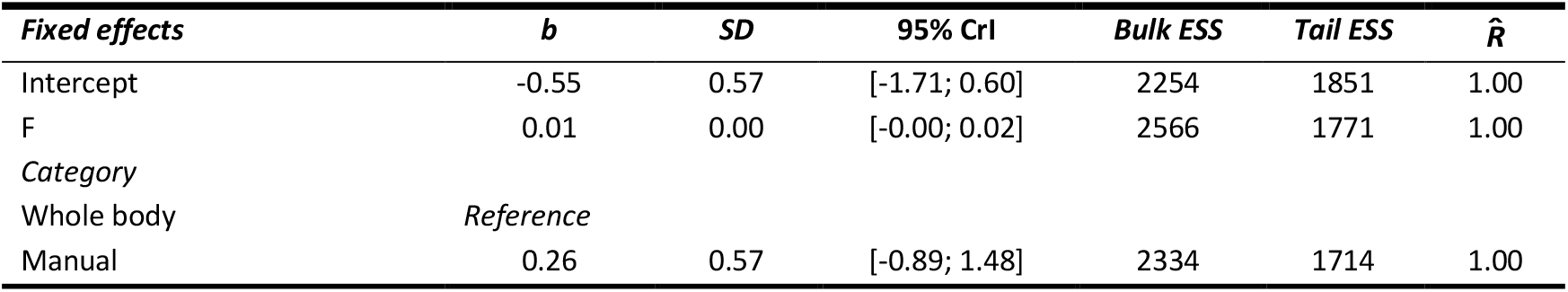
Summary of the Bayesian mixed model analysis results for the Zipf-model which included only a median value per gesture type as response variable, the frequency of gesture type as predictor and category of gesture type as control, considering only the gestures performed by Duane (*N*=15).

