## Supplemental Materials for "Variable expression of linguistic laws in ape gesture: a case study from chimpanzee sexual solicitation"

#### Supporting information 1 – Gesture types definitions

**Table S1. Ethogram of the 26 gesture types recorded in the dataset. Definitions are taken from (Hobaiter & Byrne, 2011a) and (Nishida, 2010). Video examples and illustrations of these gestures are available at www.greatapedictionary.com**

| Gesture | Description | Type |
|---|---|---|
| Beckon | Hand is moved in an upwards sweep from the elbow or wrist towards signaller. | Manual |
| Big loud scratch | Loud exaggerated scratching movement on the signaller's own body. | Manual |
| Drum | Short hard audible contact of alternate palms against an object, usually tree roots. | Manual |
| Hit object/ground | Movement of whole arm, with short hard audible contact of closed fist to an object or the ground. Includes gestures performed with one and both arms. | Manual |
| Hit object/ground with object | As 'hit object/ground' but the signaller holds an object in the hand/hands, which contacts the ground. | Manual |
| Jump | While bipedal, both feet leave the ground simultaneously, accompanied by horizontal displacement through the air. | Body |
| Leaf clipping | Strips are torn from a leaf (or leaves) held in the hand using the teeth; produces a conspicuous sound. | Manual |
| Locomote: bipedal | The signaller walks bipedally while standing up. | Body |
| Object move | Object is displaced in one direction, contact is maintained throughout movement. Includes gestures performed with one or both hands. | Manual |
| Object shake | Repeated back and forth movement of an object, usually stem of shrub, branch of tree or woody vine, performed with either one or both hands. | Manual |
| Present: genitals forwards | Signaller shows genitals to recipient. | Body |
| Raise arm | Raise arm and/or hand vertically in the air and direct palm to companion. | Manual |

| | | |
|---|---|---|
| Reach: palm | Arm extended to the recipient with the palm exposed. Typically held up or to the side, although very occasionally down. It is the palm or tip of the fingers that is closest to the recipient. | Manual |
| Reach: wrist | Arm extended to the recipient with the palm sheltered (fingers are curled), and it is either the wrist, or the back of the fingers that is reached out to the recipient. | Manual |
| Rocking: sitting | Slight or vigorous side to side movements of the body when the signaller is sitting. | Body |
| Shake arm | Small, repeated shake (adduct or abduct) of horizontally held arm at another. Includes gestures performed with either one or both arms. | Manual |
| Shake head | Small repeated back and forth motion of the head. | Manual |
| Stomp 2-feet object/ground | As 'stomp object/ground' but performed with both feet. | Manual |
| Stomp object/ground | Sole of the foot is lifted vertically and brought into a short hard audible contact with the surface being stood upon (e.g., ground, branch). | Manual |
| Stomping object/ground | As 'stomp object/ground' but performed repeatedly. | Manual |
| Swing | Large back and forth movement of the arm held below the shoulder, or of leg from the hip. Includes gestures performed with one and two arms. | Manual |
| Swing: directed | As 'swing' but the direction of the swing indicates the direction of desired movement, immediately followed by the recipient moving as indicated. | Manual |
| Swing: with object | As 'swing' but the signaller holds an object in their hand/hands (e.g., branch, leaves, etc). | Manual |
| Throw object | Object is moved and released so that there is displacement through the air after the moment of release. | Manual |
| Thrust | Rhythmic back and forth movements of the pelvis. | Body |
| Wave | Large repeated back and forth movement of the arm raised above the shoulder. | Manual |

#### Supporting information 2 - Duration distribution analysis

Before performing the GLMM analysis we analysed the distribution of the gesture duration data by (1) visually inspecting its empirical density and cumulative distribution (Figure S4) and (2) assessing its skewness and kurtosis via the visual inspection of the Cullen and Frey graph (Figure S5). Figure S5 shows the data are skewed towards low values, as almost half of the data lays between 0 and 3 seconds. Further, we fitted three theoretical distributions to the data – namely Weibull, Gamma, and Lognormal – and compared loglikelihood values (Table S2). We then plotted the three distributions and visually inspected the Q-Q, P-P, and histogram density plots (Figure S6). Finally, we compared Weibull, Gamma, and Lognormal distributions against gesture duration data distribution via goodness-of-fit tests and goodness-of-fit information criterion (Table S3 and S4), which helped identify the lognormal distribution as the best fitting one. Therefore, we proceeded with log-transforming the duration variable to best fit model assumptions.

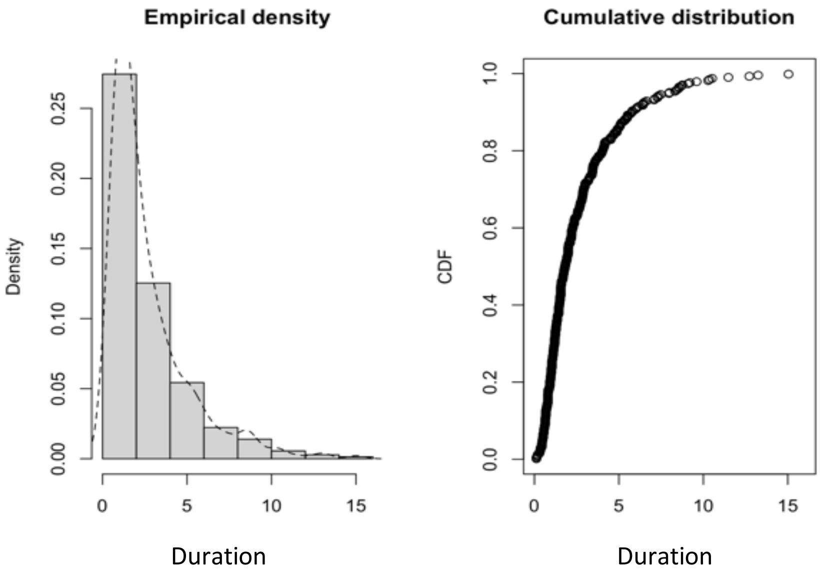

**Figure S1. Empirical distribution of gesture duration.**

Histogram and empirical cumulative distribution function (CDF) plots representing the distribution of gesture duration. Histogram bars represent sample distribution, dashed line indicates empirical density.

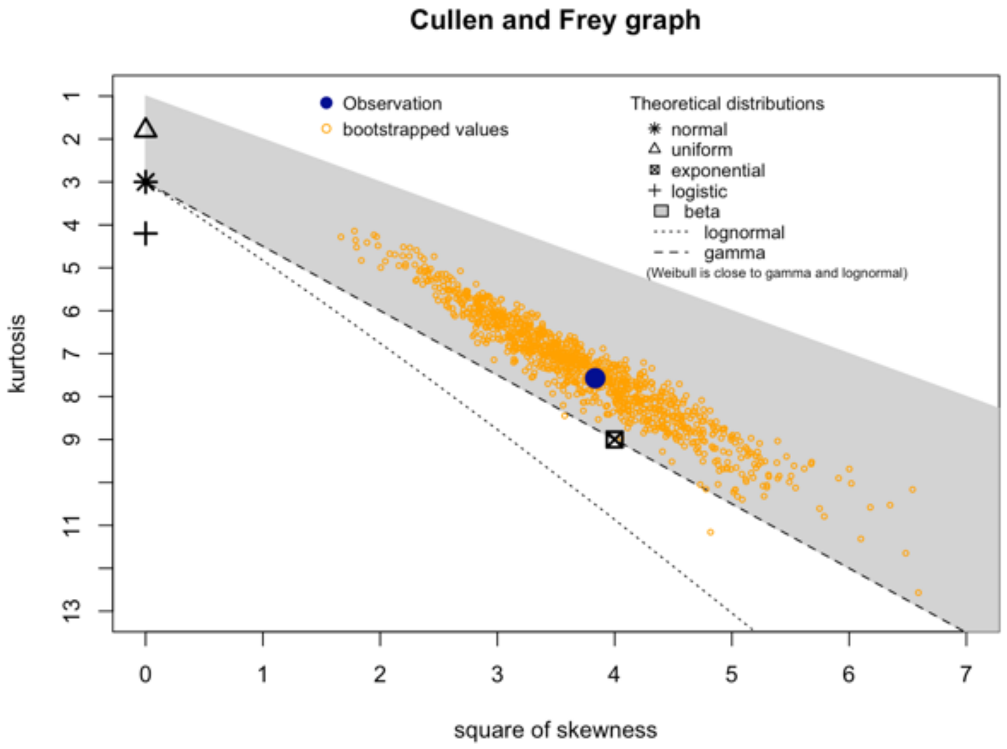

**Figure S2. Cullen and Frey graph for gesture duration.** The graph depicts the distribution of the skewness and kurtosis of gesture duration data with bootstrapped values, plotted against other theoretical distributions, namely normal, uniform, exponential, logistic, beta, lognormal, and gamma.

**Table S2. Estimate and standard error for fitting the parameters of three theoretical distributions to the distribution of the gesture duration data.**

| Distribution | Parameters | Estimate | Std Error | Loglikelihood |
|---|---|---|---|---|
| Weibull | Shape | 1.229711 | 0.04786805 | -695.5846 |
| | Scale | 2.848181 | 0.12958597 | |
| Gamma | Shape | 1.5695397 | 0.10700744 | -688.7789 |
| | Rate | 0.5933806 | 0.04755517 | |
| Lognormal | meanlog | 0.6214851 | 0.04530977 | -677.7389 |
| | sdlog | 0.8584975 | 0.03203865 | |

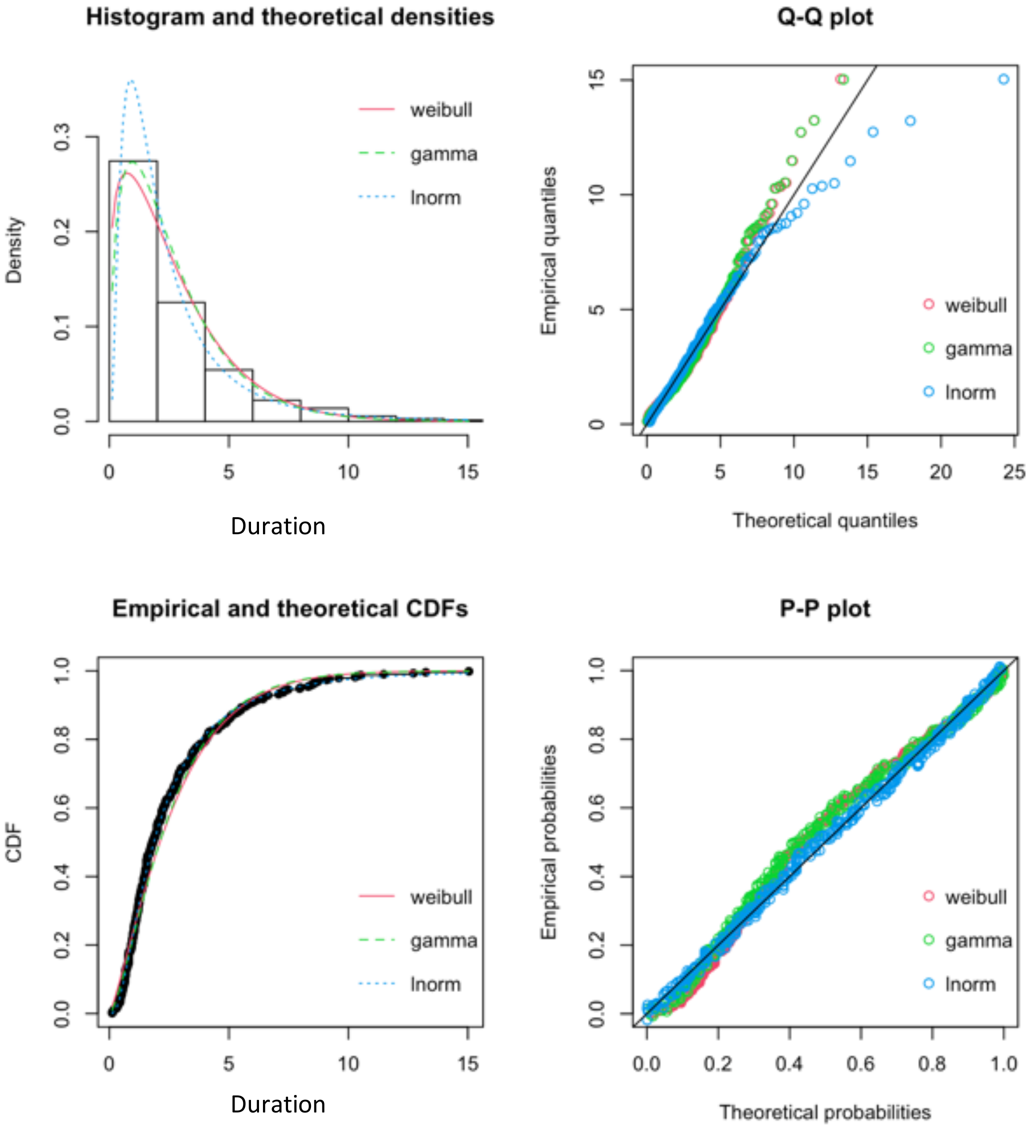

**Figure S3. Histogram and theoretical densities, Q-Q and P-P plots depicting the gesture duration data distribution against the fitted Weibull, Gamma, and Lognormal distributions.**

Histogram represents the distribution of duration data while the red, dashed green, and dashed blue lines indicate the theoretical Weibull, Gamma, and Lognormal distributions, respectively.

**Table S3. Goodness-of-fit statistics compared across fitted distributions to the gesture duration data.**

| Goodness-of-fit statistics | Weibull | Gamma | Lognormal |
|---|---|---|---|

| | | | |
|---|---|---|---|
| Kolmogorov-Smirnov statistic | 0. 07621865 | 0.08155393 | 0. 03242513 |
| Cramer-von Mises statistic | 0.62478933 | 0.55850079 | 0.03735280 |
| Anderson-Darling statistic | 3.88457421 | 3.08240994 | 0.30076490 |

**Table S4. Goodness-of-fit information criteria compared across fitted distributions.**

| Goodness-of-fit criteria | Weibull | Gamma | Lognormal |
|---|---|---|---|
| Akaike's Information Criterion | 1395.169 | 1381.558 | 1359.478 |
| Bayesian Information Criterion | 1402.936 | 1389.324 | 1367.244 |

### Supporting information 3 - Model results

**Table S5. Summary of the Bayesian mixed model analysis results for the Zipf-model which included all the data (*N*=560).**

| *Fixed effects* | *b* | *SD* | 95% CrI | *Bulk ESS* | *Tail ESS* | $\hat{R}$ |
|---|---|---|---|---|---|---|
| Intercept | 0.02 | 0.36 | [-0.67; 0.72] | 1257 | 1528 | 1.00 |
| P | 0.90 | 1.26 | [-1.25; 3.81] | 1298 | 1173 | 1.00 |
| *Category* | | | | | | |
| Whole body | *Reference* | | | | | |
| Manual | -0.24 | 0.39 | [-1.04; 0.49] | 966 | 1762 | 1.00 |
| *Random effects* | | | | | | |
| Gesture, *N*=26 | 0.80 | 0.14 | [0.57; 1.13] | 809 | 1275 | 1.00 |
| Sequence ID, *N*=377 | 0.10 | 0.07 | [0.01; 0.25] | 459 | 960 | 1.00 |
| Signaller ID, *N*=16 | 0.20 | 0.06 | [0.11; 0.34] | 1749 | 1936 | 1.00 |

**Table S6. Summary of the Bayesian mixed model analysis results for the Zipf-model which included only Duane's data (*N*=290).**

| | | | | | | |
|---|---|---|---|---|---|---|
| Intercept | -0.48 | 0.50 | [-1.46; 0.55] | 1867 | 1715 | 1.00 |
| P | 1.79 | 1.15 | [-0.39; 4.00] | 1029 | 1547 | 1.00 |
| *Category* | | | | | | |
| Whole body | *Reference* | | | | | |
| Manual | 0.31 | 0.50 | [-0.72; 1.30] | 1663 | 1672 | 1.00 |
| *Date* | | | | | | |
| 03/02/2008 | *Reference* | | | | | |
| 05/01/2008 | -0.20 | 0.11 | [-0.43; 0.02] | 3183 | 2572 | 1.00 |
| 20/01/2008 | 0.12 | 0.11 | [-0.09; 0.35] | 2765 | 2161 | 1.00 |
| *Random effects* | | | | | | |
| Gesture, *N*=15 | 0.49 | 0.17 | [0.23; 0.89] | 867 | 1541 | 1.00 |
| Sequence ID, *N*=181 | 0.11 | 0.07 | [0.01; 0.26] | 775 | 1328 | 1.00 |

**Table S7. Summary of the Bayesian mixed model analysis results for the Zipf-model which included data from all individuals but Duane (*N*=270).**

| *Fixed effects* | *b* | *SD* | 95% CrI | *Bulk ESS* | *Tail ESS* | $\hat{R}$ |
|---|---|---|---|---|---|---|
| Intercept | 0.27 | 0.46 | [-0.61; 1.21] | 1442 | 1499 | 1.00 |
| P | 0.00 | 0.00 | [-0.00; 0.01] | 2428 | 1993 | 1.00 |
| *Category* | | | | | | |
| Whole body | *Reference* | | | | | |
| Manual | -0.36 | 0.50 | [-1.38; 63] | 1565 | 1769 | 1.00 |
| *Random effects* | | | | | | |
| Gesture, *N*=18 | 0.87 | 0.19 | [0.58; 1.32] | 1194 | 1657 | 1.00 |
| Sequence ID, *N*=196 | 0.20 | 0.10 | [0.01; 0.38] | 468 | 1053 | 1.01 |

**Table S8. Summary of the Bayesian mixed model analysis results for the Menzerath-model which included all the data (*N*=530).**

| *Fixed effects* | *b* | *SD* | 95% CrI | *Bulk ESS* | *Tail ESS* | $\hat{R}$ |
|---|---|---|---|---|---|---|
| Intercept | 0.69 | 0.14 | [0.41; 0.97] | 1316 | 1809 | 1.00 |
| **Sequence Size** | **-0.18** | **0.04** | **[-0.26; -0.11]** | **2374** | **1915** | **1.00** |
| PWB | -0.31 | 0.20 | [-0.71; 0.08] | 2795 | 2529 | 1.00 |
| *Random effects* | | | | | | |
| Signaller ID, N=16 | 0.36 | 0.11 | [0.18; 0.62] | 938 | 1501 | 1.00 |
| Sequence ID, N=359 | 0.31 | 0.10 | [0.07; 0.47] | 356 | 539 | 1.01 |

**Table S9. Summary of the Bayesian mixed model analysis results for the Menzerath-model which included only Duane's data (*N*=273).**

| *Fixed effects* | *b* | *SD* | 95% CrI | *Bulk ESS* | *Tail ESS* | $\hat{R}$ |
|---|---|---|---|---|---|---|
| Intercept | 0.92 | 0.13 | [0.67; 1.17] | 4365 | 2026 | 1.00 |
| **Sequence Size** | **-0.23** | **0.04** | **[-0.31; -0.15]** | **4074** | **2145** | **1.00** |
| PWB | -0.69 | 0.58 | [-1.89; 0.38] | 4900 | 2168 | 1.00 |
| *Date* | | | | | | |
| 03/02/2008 | *Reference* | | | | | |
| 05/01/2008 | 0.06 | 0.14 | [-0.22; 0.34] | 4821 | 2130 | 1.00 |
| 20/01/2008 | 0.48 | 0.13 | [0.22; 0.75] | 5325 | 2610 | 1.00 |
| *Random effects* | | | | | | |
| Sequence ID, N=181 | 0.11 | 0.08 | [0.00; 0.30] | 995 | 1009 | 1.00 |

**Table S10. Summary of the Bayesian mixed model analysis results for the Menzerath-model which included data from all individuals but Duane (*N*=257).**

| *Fixed effects* | *b* | *SD* | 95% CrI | *Bulk ESS* | *Tail ESS* | $\hat{R}$ |
|---|---|---|---|---|---|---|
| Intercept | 0.33 | 0.18 | [-0.04; 0.70] | 2167 | 2241 | 1.00 |
| Sequence Size | 0.01 | 0.08 | [-0.14; 0.16] | 2866 | 2438 | 1.00 |
| PWB | -0.24 | 0.22 | [-0.69; 18] | 3474 | 2226 | 1.00 |
| *Random effects* | | | | | | |
| Sequence ID, *N*=187 | 0.42 | 0.11 | [0.16; 0.62] | 466 | 423 | 1.01 |
| Signaller ID, *N*=15 | 0.37 | 0.14 | [0.14; 0.70] | 842 | 902 | 1.00 |

**Abbreviations**: *b*= Estimated mean of the posterior distribution; *SD*= Standard deviation of the posterior distribution; *CrI*= Two-sided 95% Credible intervals based on quantiles; *Bulk ESS*= the effective sample size for rank normalized values using split chains; *Tail ESS*= the minimum of the effective sample sizes for 5% and 95% quantiles; $\hat{R}$=R hat value, provides information about the convergence of the Bayesian model algorithm.

#### Supporting information 4

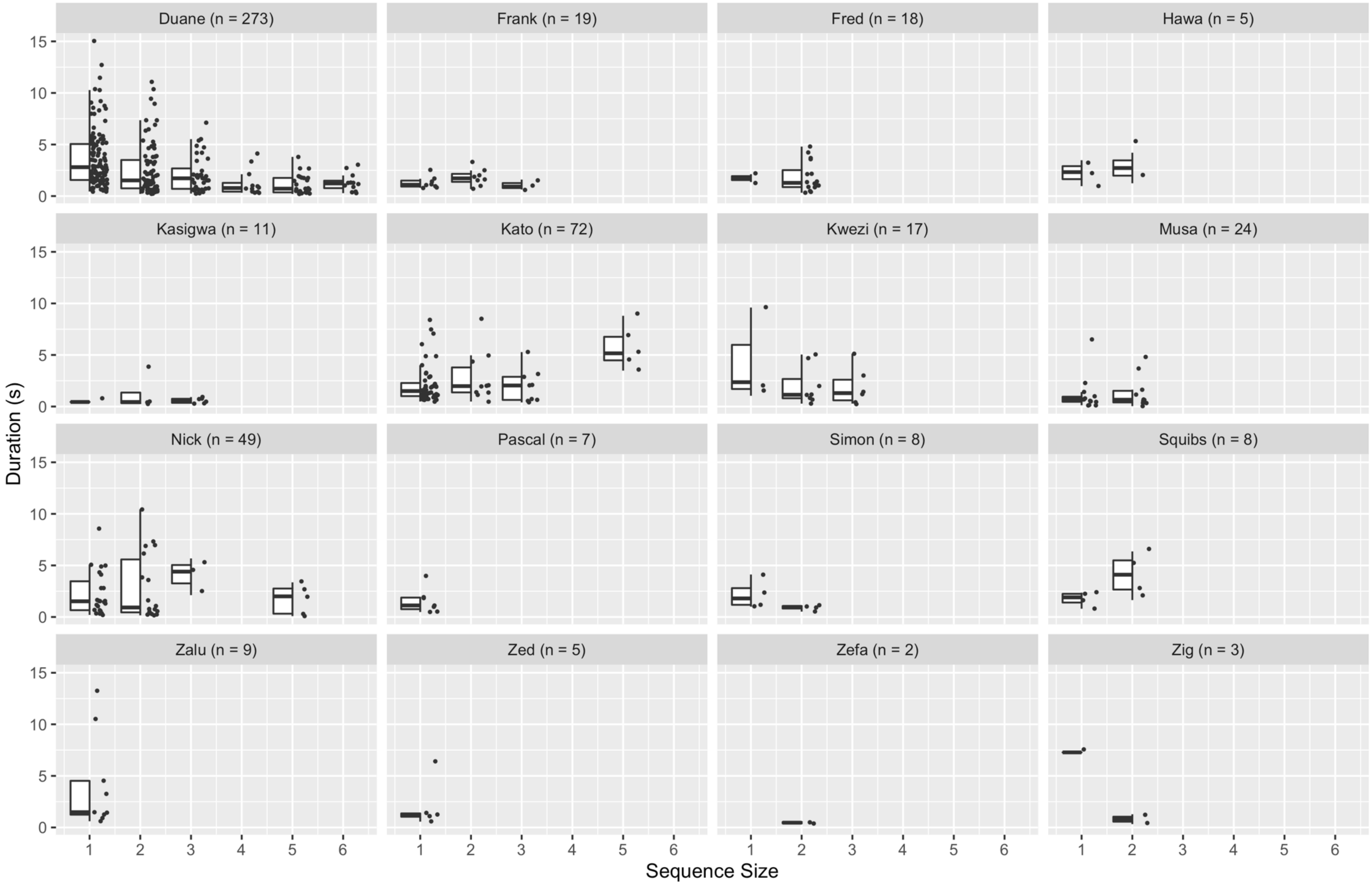

**Figure S4. Distribution of gesture durations based on sequence size for each of the 16 individuals in the dataset.** Points represent individual gesture tokens. Boxplots show median (black central bar), interquartile range (boxes), maximum and minimum values exploding outliers (whiskers). n indicates sample size for each individual.

#### Supporting information 5 – Visual inspection of sequence structure

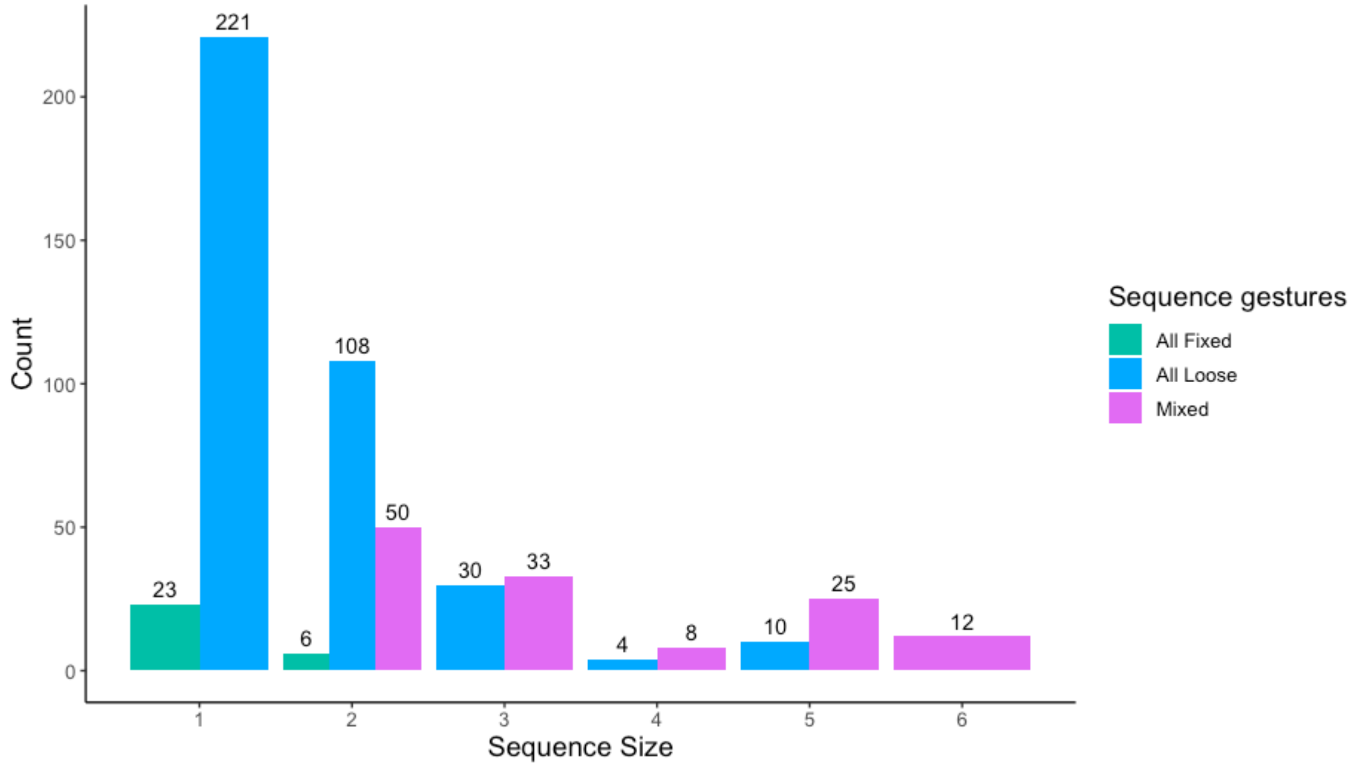

**Figure S5. Bar chart showing the frequency distribution of the three different types of sequences depending on their sequence size.**

Green sequences comprise only *fixed* duration gestures, blue sequences only *loose* duration gestures. Pink bars represent sequences formed by a mix of *loose* and *fixed* duration gestures. Numbers above bars indicate the frequency of each sequence type per sequence size.

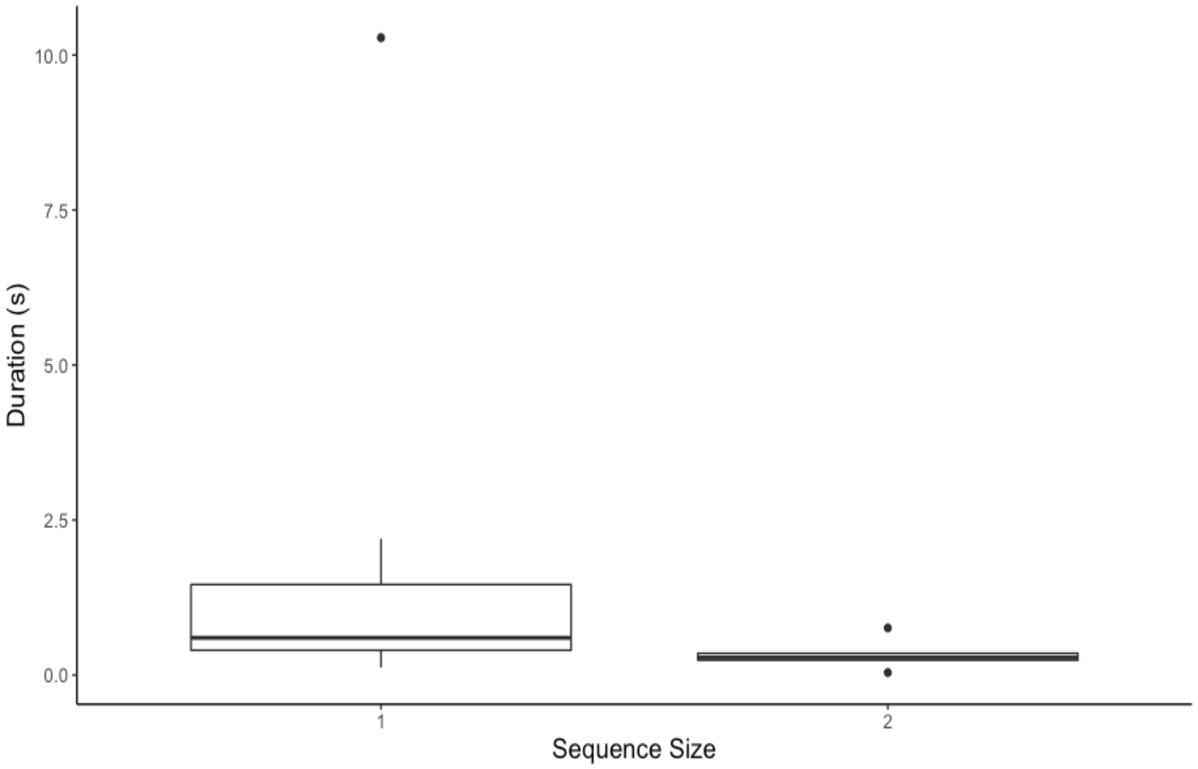

**Figure S6. Boxplots of the duration of gestures with constrained duration (i.e., *fixed* duration gestures) in sequences formed of *fixed* duration gestures solely.**

Boxplots show median (black central bar), interquartile range (boxes), maximum and minimum values exploding outliers (whiskers) and outliers (circles).

$$M = \sum_{i=1}^{N} T_i \qquad (3)$$

where $T_i$ is the total duration of the $i$th sequence and *N* is the number of sequences.

In turn,

$$T = \sum_{j=1}^{n_i} t_{ij} \qquad (4)$$

where $t_{ij}$ is the duration of the $j$th element of the $i$th sequence and $n_i$ is the size of the $i$th sequence. Given that the mean duration of gestures from the $i$th sequence can be expressed as $\left(t_{ij}\right)_i = \frac{T_i}{n_i}$, $M$ can be defined as

$$M = \sum_{i=1}^{N} n_i \left(t_{ij}\right)_i \quad (5)$$

*M* was calculated through this equation and was tested to assess whether it is significantly small. We performed a similar permutation test to that conducted to test for the significance of *L*, to check whether *M* was significantly small as compared to the values generated by random permutation of the data (Zipf, 1936). In such case, $n_i$ has the role of $p_i$ and $\left(t_{ij}\right)_i$ has the role of $e_i$ in the test, with $n_i$ and $\left(t_{ij}\right)_i$ remaining constant during the test.

The permutation test produces a left *p*-value to check if *L* (or *M*) is significantly small and a right *p*-value to check if *L* (or *M*) is significantly large compared to the distribution of the values created by a permutation of the data (Heesen et al., 2019). The total number of permutations carried out was $R=10^5$.

#### Results of one-tailed analyses

##### Zipf's law of brevity

##### Do chimpanzee sexual solicitation gestures follow Zipf's law of brevity?

We did not find a pattern in agreement with Zipf's law of brevity; there was no evidence for a significant negative correlation between mean gesture type duration (*d*) and frequency of use (*f*) (Spearman correlation: $r_s$=0.30, *n*=26, *p*=0.066), in agreement with the Bayesian model analysis. Consistent with this result, the compression test revealed that the expected mean code length of gesture types *L* had a magnitude of 2.39s and was not significantly small ($p_{left}$=0.951). Rather, *L* was significantly big ($p_{right}$=0.05, Figure S7).

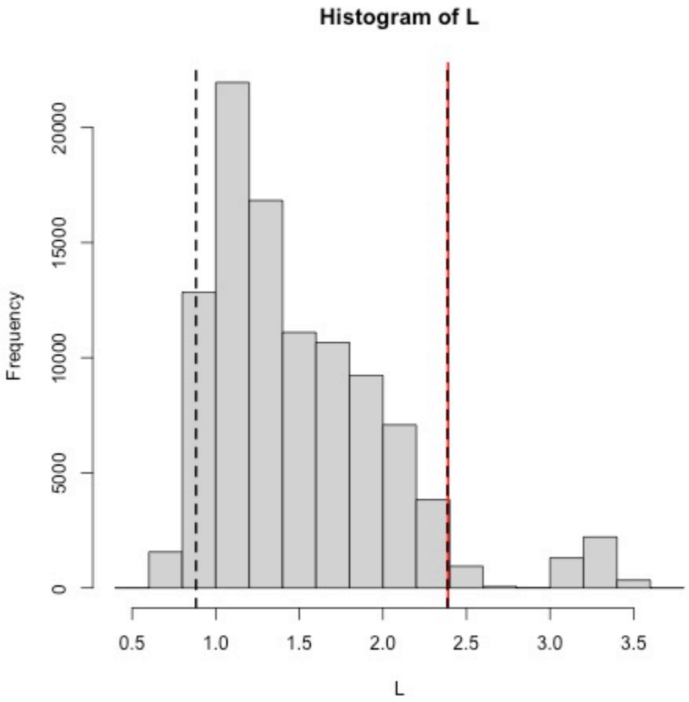

**Figure S7. Histogram showing the distribution of the permuted *L* values. Observed *L* value is highlighted with the red continuous line. Black dashed line indicates the lower and upper 5% of the permuted data.**

**Subset analysis: whole-body and manual gesture types.**

We found no evidence for a negative correlation between *d* and *f* when separating whole-body gestures from manual gestures (Spearman's rank correlation: whole-body, $r_s$=-0.3, *n*=5, *p*=0.342; manual, $r_s$ =0.42, *n*=21, $p_{left}$=0.969). Rather, manual gestures showed a significant positive correlation ($r_s$ =0.42, *n*=21, $p_{right}$=0.031). Compression tests revealed that for whole-body gestures, *L*=0.13s and was neither significantly big or small ($p_{left}$=0.174, $p_{right}$=0.817), and for manual gestures, *L*=2.26s and, if anything, tended towards being significantly big ($p_{right}$=0.058) rather than small ($p_{left}$=0.942; Figure S8).

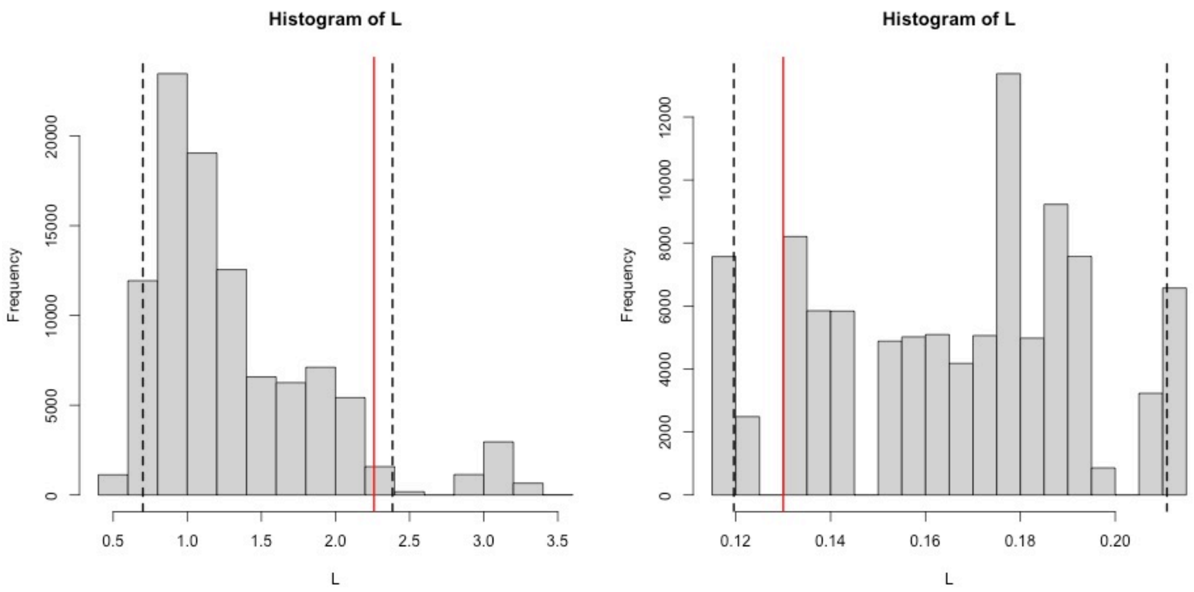

**Figure S8. Histograms showing the distribution of the permuted *L* values for manual gestures (left) and whole-body gestures (right). Observed *L* value is highlighted with the red continuous line. Black dashed lines indicate the upper and lower 5% of the permuted data***.*

**Do chimpanzee sexual solicitation gesture sequences follow Menzerath's law?**

We tested Menzerath's law in 359 sequences, composed of 530 gesture tokens; there was no evidence for a negative relationship between mean constituent duration and sequence size (Spearman's rank correlation: $r_s$=-0.08 *n*=359, *p*=0.076). However, the compression test revealed that the total sum of the duration of each sequence *M* had a value of 1300.67 and was significantly small (*n*=359, *p*=0.003; Figure S9).

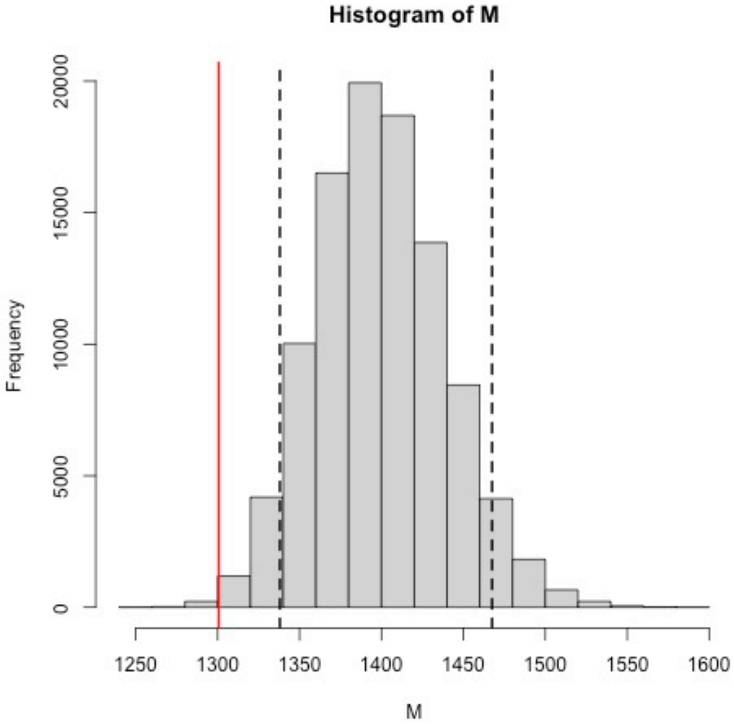

**Figure S9. Histogram showing the distribution of the permuted *M* values. Observed *M* value is highlighted with the red continuous line. Black dashed line indicates lower 5% of the permuted data.**

#### Discussion

The results from the correlation analysis must be taken with caution as this analysis does not control for individual variation, gesture type, and sequence in which the gesture is performed. The Bayesian model that included these factors and which tested for Zipf's law of brevity (Zipf-model) was similar to the respective null-model, suggesting that frequency of gesture type within the dataset and category of gesture type did not predict gesture duration.

**Table S11. Summary of the Bayesian mixed model analysis results for the Zipf-model which included only a median value per gesture type as response variable, the frequency of gesture type as predictor and category of gesture type as control (*N*=26).**

| *Fixed effects* | *b* | *SD* | 95% CrI | *Bulk ESS* | *Tail ESS* | $\hat{R}$ |
|---|---|---|---|---|---|---|
| Intercept | 0.10 | 0.33 | [-0.55; 0.77] | 2267 | 1782 | 1.00 |
| F | 0.00 | 0.00 | [-0.00; 0.01] | 2861 | 1404 | 1.00 |
| *Category* | | | | | | |
| Whole body | *Reference* | | | | | |
| Manual | -0.24 | 0.36 | [-0.96; 0.48] | 1995 | 1776 | 1.00 |

**Table S12. Summary of the Bayesian mixed model analysis results for the Zipf-model which included only a median value per gesture type as response variable, the frequency of gesture type as predictor and category of gesture type as control, considering only the gestures performed by Duane (*N*=15).**

| *Fixed effects* | *b* | *SD* | 95% CrI | *Bulk ESS* | *Tail ESS* | $\hat{R}$ |
|---|---|---|---|---|---|---|
| Intercept | -0.55 | 0.57 | [-1.71; 0.60] | 2254 | 1851 | 1.00 |
| F | 0.01 | 0.00 | [-0.00; 0.02] | 2566 | 1771 | 1.00 |
| *Category* | | | | | | |
| Whole body | *Reference* | | | | | |
| Manual | 0.26 | 0.57 | [-0.89; 1.48] | 2334 | 1714 | 1.00 |
